# Reductive stress contributes to neuronal injury in ischemic stroke: fructose protection via the reverse TCA cycle

**DOI:** 10.64898/2026.11.26.769592

**Authors:** T. Ren, Y. Feng, J. Liang, X. Song, Y. Wang, X. Ji, B. Zhou

**Author notes:** Corresponding author. (B.Zhou).

## Abstract

Acute ischemic stroke presents a therapeutic paradox: reperfusion is essential for neuroprotection but triggers oxidative injury. The limited efficacy of antioxidant therapies suggests that critical upstream mechanisms remain unresolved. Here, we identified marked NADH accumulation in ischemic brain lesions, revealing a state of reductive stress that disrupts metabolism and impairs mitochondrial function. Targeting this reductive stress provides earlier neuroprotection than antioxidant approaches, uncovering the therapeutic potential of upstream metabolic regulation. We found that fructose induces a non-canonical TCA cycle reversal via fructose-1-phosphate (F1P), decreases the NADH/NAD□ ratio, diminishes mitochondrial damage, and protects neurons against ischemic injury. This metabolic rewiring also replenishes key biosynthetic intermediates, sustaining cellular viability. Consistently, clinical evidence supports the neuroprotective potential of fructose, suggesting a novel therapeutic strategy for acute ischemic stroke.

## Introduction

The brain is highly vulnerable to ischemia due to its high energy demand, limited metabolic reserve, and reliance on oxidative phosphorylation of neurons (*1–4*), making early metabolic intervention critical for stroke treatment. Although ischemia-induced metabolites have been proposed to contribute to reperfusion-associated free radical bursts (*5*), stroke research has largely focused on oxidative stress arising after reperfusion (*6–12*). In contrast, the metabolic status of reductive imbalance during the sole ischemic phase remains largely unexplored.

During energy metabolism, electrons are extracted through glycolysis and the TCA cycle, carried by reducing coenzymes to the mitochondrial respiratory chain, where oxygen serves as the terminal electron acceptor. Under physiological conditions, electrons cycle between redox couples, including NAD□/NADH pair, enabling continuous NAD□ regeneration and oxidative phosphorylation. Hypoxia disrupts this electron flux, theoretically causing reducing equivalents to accumulate and giving rise to reductive stress. However, whether such metabolic stress occurs in ischemic brain tissue, which redox carriers are affected, and whether this state represents a therapeutic opportunity remain unknown. Here, we show that initial ischemic injury is driven by excessive electron and NADH accumulation, rather than early ROS elevation. Mass spectrometry imaging and quantitative analysis confirmed that NADH levels reach approximately 240 µM/mg in ischemic brain tissue, which triggers neuronal mitochondrial and endoplasmic reticulum stress, TCA cycle dysregulation, and subsequent mitochondrial damage.

In searching for metabolic interventions during ischemia, we unexpectedly found that ischemic preconditioning markedly elevated cerebral fructose levels, consistent with previous work linking fructose-driven glycolysis to extreme hypoxia tolerance in naked mole-rats (*13*). Fructose is transported into neurons via GLUT5 and catabolized by KHK to generate fructose-1-phosphate (F1P), which serves as a signal to redirect pyruvate into the reversed TCA cycle via pyruvate carboxylase (PC). This pathway oxidizes excess NADH back to NAD□, restores redox homeostasis, and replenishes metabolic intermediates. Fructose effectively alleviates acute ischemic brain injury, exerts potent neuroprotection in aged and diabetic comorbidity mice, and improves NIHSS score in our clinical trial, highlighting its promising translational potential.

### Ischemic NADH accumulation triggers neuronal injury

To define metabolic alterations during acute ischemia, we performed MALDI-TOF mass spectrometry imaging on ischemic brain tissue. Unexpectedly, most metabolic intermediates remained abundant rather than being depleted within ischemic lesions (Fig. 1A, Fig. S1A). By contrast, NADH accumulated dramatically (Fig. 1A). Absolute quantification revealed that tissue NADH levels increased from 199 µM/µg at 1 h to 243 µM/µg at 2 h post ischemia (fig. S1B-S1D), indicating that NADH acts as the primary electron carrier during ischemia, rather than GSH and NADPH, which exhibit distinct temporal dynamic patterns (fig. S1E).

**Fig. 1.**
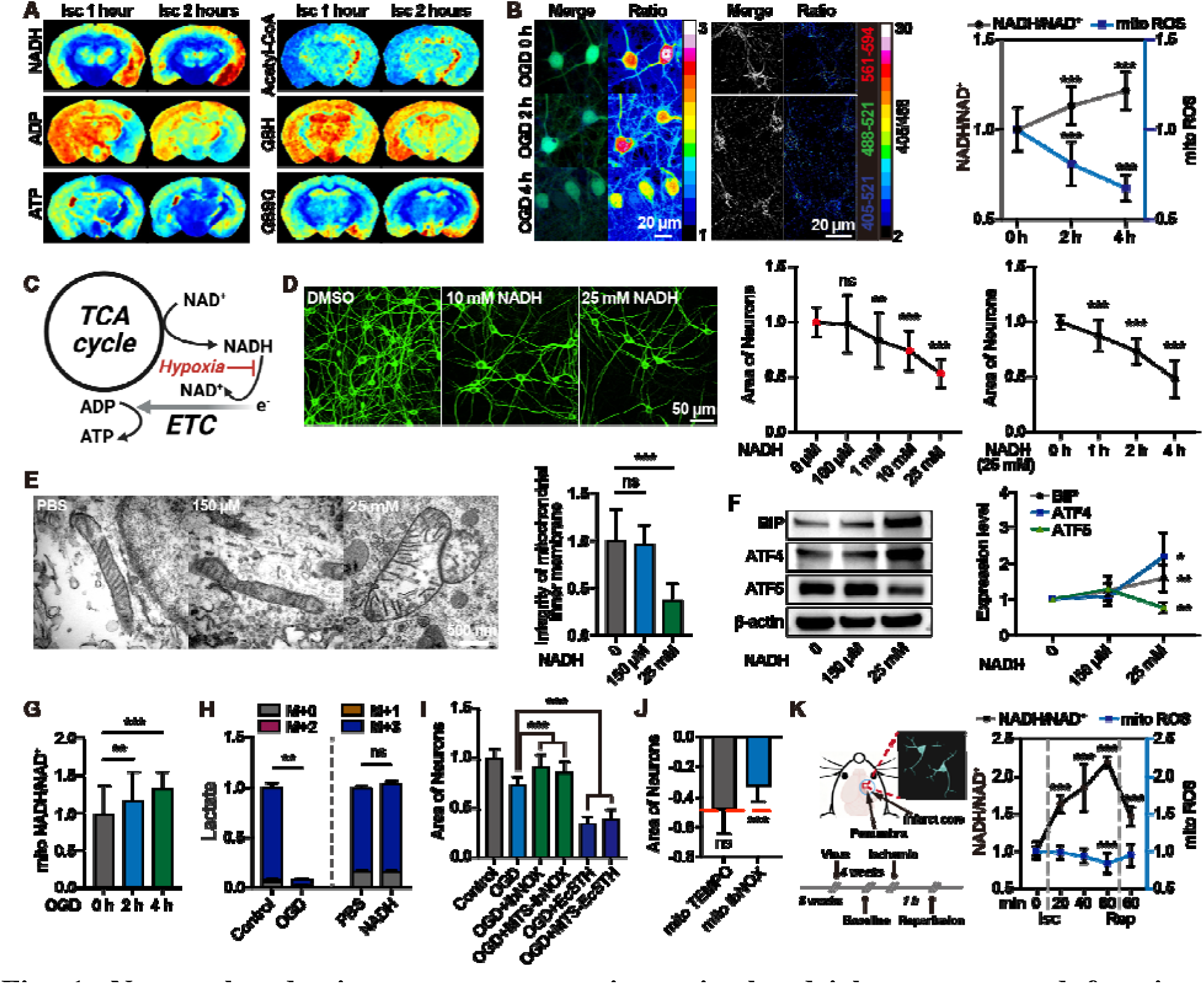
Neuronal reductive stress compromises mitochondrial structure and function, driving global disruption of cellular redox homeostasis. **(A)** Mass spectrometry imaging revealing dynamic shifts in metabolite abundance across sequential ischemia and reperfusion time points. **(B)** Quantification of cellular NAD□/NADH and mitochondrial ROS levels in primary neurons subjected to 2 h or 4 h OGD via the SoNar and mitoROS biosensors, respectively (N=5, n>6). **(C)** Schematic illustrating OGD-mediated electron transport chain inhibition, which uncouple de novo NADH production from its respiratory consumption. **(D)** Dose- and time-dependent loss of viability in primary neurons following exogenous NADH supplementation (N=5, n>6). **(E)** Transmission electron microscopy (TEM) and quantitative analysis of inner mitochondrial membrane integrity in primary neurons incubated with either 150 µM or 25 mM NADH (N=6, n>10). **(F)** Western blot profiling of mitochondrial and endoplasmic reticulum (ER) stress signaling cascades in primary neurons treated with 150 µM or 25 mM NADH for 4 h (N>4). **(G)** Time-course quantification of intramitochondrial NADH/NAD□ ratios across progressive OGD stages using the mitochondrially targeted MTS-SoNar biosensor (N=5, n>6). **(H)** Isotope-labeled glucose tracing to quantify lactate production in primary neurons under distinct experimental conditions (N=3 to 4). **(I)** Genetic manipulation of intracellular NAD(H) pools via overexpression of lbNOX, MTS-lbNOX, EcSTH, or MTS-EcSTH in primary neurons. MAP2 immunofluorescence staining was subsequently performed to assess neuronal survival and morphological preservation after OGD injury (N=5, n>10). **(J)** Quantification of neuronal survival in primary cultures transfected with empty vector or mito-lbNOX, subjected to OGD-reperfusion, and co-treated with vehicle DMSO or mitoTEMPO during OGD exposure (N=5, n>5). **(K)** In vivo two-photon imaging of cortical neurons in C57BL/6J mice transduced with brain-targeted SoNar and mitoROS biosensors. Serial signal acquisition was conducted prior to ischemia, during tMCAO occlusion, and following reperfusion to quantify dynamic redox shifts within tissue within the injured lesion (N=4, n>=3). Data are presented as means ± SD. Multiple-group comparisons were performed using one-way analysis of variance (ANOVA), followed by Dunnett’s post-hoc test (B, D, E, F, G, I and L); comparisons between two groups were conducted using the unpaired Student’s t-test (H and J). *P < 0.05, **P < 0.01, ***P < 0.001; ns, not significant.

As NADH levels exhibited a progressively rising trend during ischemia, and the TCA cycle serves as the major site of NADH production with adequate substrate such as acetyl-CoA (Fig. 1A), we hypothesized that the TCA cycle continues despite oxygen deprivation, while respiratory chain suppression blocks NADH reoxidation to NAD□ and its utilization for ATP production (Fig. 1A), indicating ischemia uncouples TCA and ETC at the NADH node (Fig. 1C). To determine whether NADH accumulation directly contributes to brain cells injury, we elevated NADH by culture supplement. In mixed neuron-astrocyte cultures, adding 25 mM NADH to the culture medium increased intracellular NADH to approximately twice the level (500 µM/µg) that was measured in ischemic brain tissue (fig. S1F). Notably, NADH overload selectively damaged neurons over astrocytes (fig. S1G, S1H), despite comparable increases of the NADH/NAD□ ratio (fig. S1K, S1L), which is consistent with the greater neuronal vulnerability observed under oxygen-glucose deprivation (OGD) (fig. S1I, S1J). We further performed dose assessment and found that 1 mM NADH triggers, while 25 mM NADH promotes, substantial neuronal death within 4 h (Fig. 1D), which is correlated with elevated NADH/NAD□ (fig. S1M, S1N). To monitor redox dynamics directly, we expressed the genetically encoded NADH/NAD□ sensor SoNar and the mitochondrial ROS reporter in neurons (*14, 15*) both in vitro and in vivo (Fig. 1B, 1K), and revealed a progressive increase in the NADH/NAD□ ratio throughout ischemia, accompanied by a decline in mitochondrial ROS production. Thus, acute ischemia drives neurons into a highly reduced state that triggers reductive stress and subsequent neuronal injury (*16*).

We next investigated the subcellular effects of NADH overload. Although NADH induced both mitochondrial and ER stress (Fig. 1F), it selectively disrupted mitochondrial morphology with minimal effects on the Golgi and ER in cultured cortical neurons (Fig. 1E, fig. S2A-S2D) and ischemic lesions (fig. S2E, S2F). To link this mitochondrial vulnerability to ischemic injury, we found that the mitochondrial matrix NADH/NAD□ ratio increased upon OGD (Fig. 1G). Similarly, respiratory complex I inhibition also caused greater neuronal injury than in astrocytes (fig. S2G-S2H). Unlike tumor cells, neurons cannot compensate through glycolytic fermentation: stable isotope tracing showed that neither OGD nor NADH supplementation increased lactate production (Fig. 1H), suggesting that neurons fail to regenerate NAD□ through compensatory glycolytic fermentation as an alternative route, which was confirmed by exacerbated injury from oxygen deprivation (fig. S2K).

Since neurons are unlikely to counteract NADH reductive stress by fermentation, we then asked whether NAD(H) regulation determines neuronal survival. We genetically manipulated cellular NAD(H) by overexpressing lbNOX (NADH oxidase) (*17–19*), EcSTH (NADH-producing enzyme) (*20*), and their mitochondrial variants (fig. S2J). lbNOX or MTS-lbNOX improved neuronal survival after OGD, while EcSTH/MTS-EcSTH exacerbated cell death (Fig. 1I). These results demonstrate that neuronal survival is indiscernibly governed by compartment and total NAD(H) homeostasis. Interestingly, we observed that NADH overload induced mitochondrial DNA (mtDNA) release into the cytoplasm (fig. S2L, S2M), a marker of severe mitochondrial damage. Notably, mitochondrial DNA leakage may contribute to adverse inflammatory responses (*21*). To clarify the mechanism by which NADH disrupts mitochondrial membrane structure, we selectively elevated mitochondrial NADH levels via MTS-EcSTH. We found that MTS-EcSTH treatment reduced cardiolipin content (fig. S2N, S2O), decreased inner membrane fluidity (fig. S2P, S2Q), and increased mitochondrial permeability (fig. S2R). The results suggest that reductive stress destabilizes the mitochondrial membrane structure, thereby inducing mitochondrial damage. To further dissect the predominant stress driving ischemic injury, we quantified mitochondrial cardiolipin levels under distinct interventions. We found that, in contrast to mitoTEMPO (which primarily suppresses ROS) (*22, 23*), only mitochondrial LbNOX restored cardiolipin abundance and rescued neuronal viability under OGD (Fig. 1J, Fig. S2S), demonstrating that reductive stress is the dominant driver of neuronal damage during the ischemic phase.

### Electron overload underlies reductive stress

Given that glucose is the principal energy substrate of the brain, and the respiratory chain is required to transfer electrons derived from its catabolism to oxygen (Fig. 2A). We first measured metabolites related to glycolysis at different time points after ischemia (Fig. 2B). The results revealed distinct temporal changes in glycerol-3-phosphate (G3P) and 1,3-bisphosphoglycerate (1,3-BPG) metabolites (the only enzymatic reactions are coupled to electron transfer in glycolysis). At 1-hour post-ischemia, G3P was largely converted to 1,3-BPG. By contrast, massive G3P accumulation and markedly reduced 1,3-BPG production were detected at 2 hours post-ischemia, indicating that the enzymatic reaction tended to drive the reduction of 1,3-BPG back to G3P, likely because prolonged ischemia drives glycolysis toward a highly reduced state in which electron disposal becomes limiting. In addition, multiple enzymatic reactions in TCA cycle involve electron transfer (Fig. 2B). We found that dehydrogenation reactions catalyzed by IDH, KGDHC, and SDH still proceeded at 1-hour post-ischemia, indicating that endogenous electron acceptors could buffer excess electrons, which was consistent with the dehydrogenation during glycolysis (Fig. 2B). Nevertheless, all these dehydrogenation reactions slowed down at 2 hours post-ischemia, and some even reversed their direction, indicating that cells consequently engage alternative electron acceptors such as fumarate (*24*) and succinyl-CoA. Notably, succinyl-CoA showed greater capacity for reverse electron transfer under conditions of electron accumulation, indicating its superior electron-accepting potential. Because NAD(H) mediates electron transfer across numerous enzymatic reactions, we hypothesized that electron overload disrupts metabolism through NADH over-reduction (Fig. 2C).

**Fig. 2.**
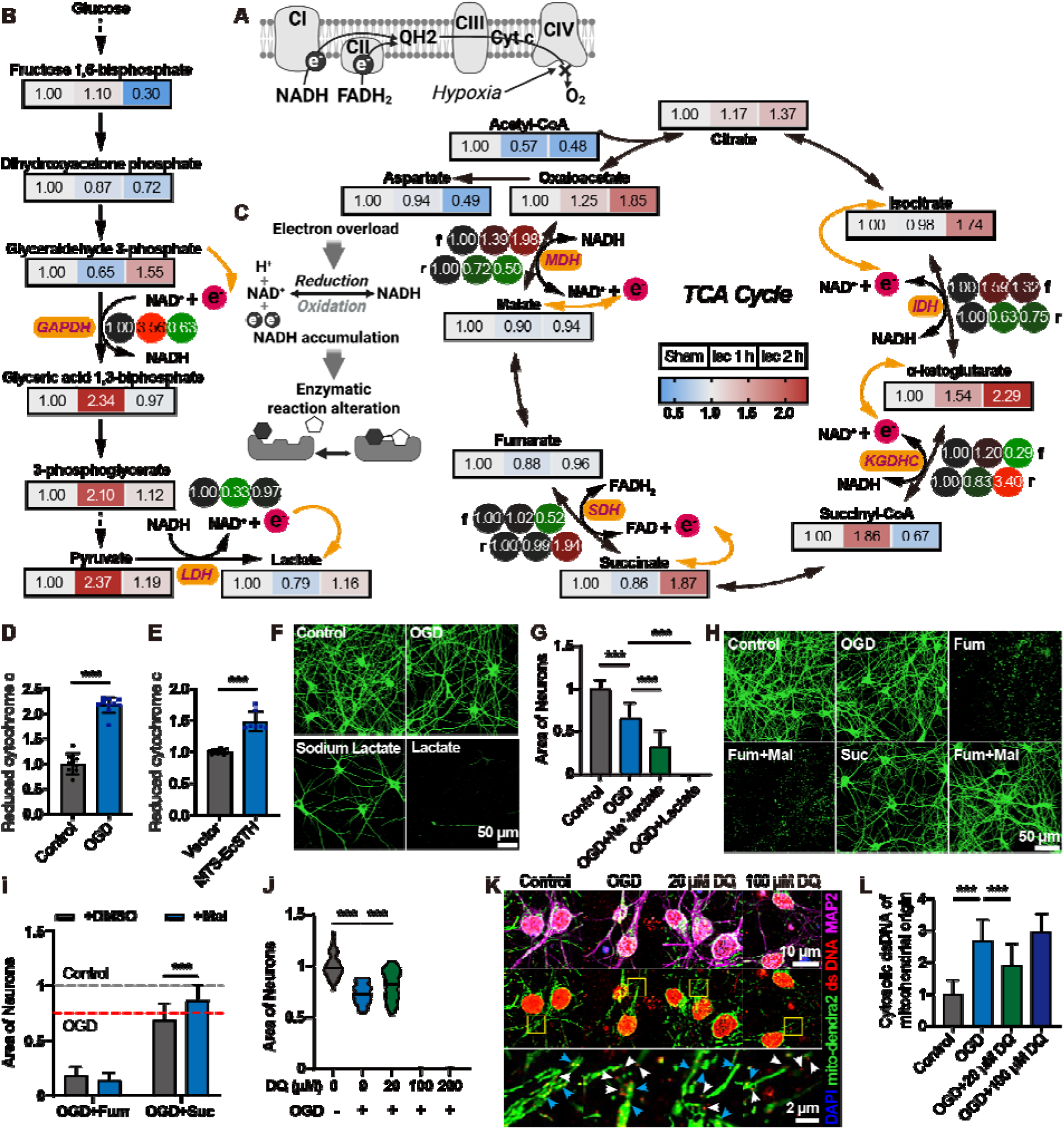
Excessive electron accumulation constitutes the core driver of reductive stress, which remodels enzymatic reactions by perturbing cellular NAD(H) pools. **(A)** Schematic illustrating the shortage of terminal electron acceptors under hypoxia, which blocks electron transfer from NADH and FADH via the electron transport chain. **(B)** LC-MS-based quantification of glycolytic and TCA-cycle metabolite abundances across sequential ischemic time points, followed by calculation of catalytic rates for electron-transferring enzymatic reactions (N=3). **(C)** Schematic illustrating how imbalanced interconversion between NADH and NAD□ driven by surplus electrons reshapes downstream enzymatic activity. **(D)** Quantification of reduced cytochrome c abundance in primary neurons subjected to 4 h OGD (N=8 to 10). **(E)** Measurement of reduced cytochrome c levels in primary neurons transfected with empty vector or mitochondrially targeted MTS-EcSTH (N=7 to 12). **(F and G)** Primary neurons were exposed to OGD-reperfusion injury. Lactate or sodium lactate was supplemented exclusively during OGD and washed out upon reperfusion; MAP2 immunostaining was performed to quantify neuronal survival (N=5, n>10). **(H and I)** Primary neurons exposed to OGD-reperfusion, with fumarate or succinate co-administered during OGD alongside vehicle DMSO or malonate (Mal), an SDH inhibitor. All metabolites were removed at reperfusion, and MAP2 staining was used to assess neuronal viability (N=5, n>6). **(J)** Primary neurons were exposed to OGD-reperfusion and treated with DMSO, 20 µM duroquinone (DQ), or 100 µM DQ during OGD (compounds washed out at reperfusion). Neuronal survival was quantified via MAP2 immunofluorescence (N=4, n>5). **(K and L)** Primary neurons were labeled via mito-dendra2 transduction and subjected to 4 h OGD. Cultures received DMSO, 20 µM DQ, or 100 µM DQ throughout OGD; immunofluorescence staining was performed to quantify mitochondrial DNA release (N=6, n>5). Data are presented as means ± SD. Multiple-group comparisons were performed using one-way analysis of variance (ANOVA), followed by Dunnett’s post-hoc test (G, J and L); two-way analysis of variance (ANOVA) was performed, followed by Sidak’s multiple comparisons test (I); comparisons between two groups were conducted using the unpaired Student’s t-test (H and J). *P < 0.05, **P < 0.01, ***P < 0.001; ns, not significant.

OGD induced marked accumulation of reduced cytochrome c (Fig. 2D), reflecting impaired electron consumption (Fig. 2A). Elevating mitochondrial NADH using MTS-EcSTH similarly increased reduced cytochrome c levels (Fig. 2E), demonstrating that respiratory-chain hyper-reduction occurs when NADH generation exceeds mitochondrial oxidative capacity, thereby compromising mitochondrial integrity (*5, 25*). To determine whether electron burden directly affects neuronal survival, we increased endogenous electron input using lactate, which releases electrons upon oxidation. Lactate supplementation further decreased the NAD□/NADH ratio and exacerbated OGD-induced neuronal injury (Fig. 2F, 2G; fig. S3Q), indicating that although lactate serves as a metabolic substrate, it cannot support neuronal survival during stroke because it aggravates electron overload. Similarly, succinate supplementation increased neuronal death, whereas inhibition of succinate dehydrogenase with malonate, which blocks electron release from succinate, attenuated this toxicity (Fig. 2H, 2I). Although fumarate can act as an alternative electron acceptor, exogenous fumarate unexpectedly caused extensive neuronal toxicity, suggesting that its detrimental effects may outweigh its potential electron-buffering capacity (*26*) (Fig. 2H, 2I). We next evaluated duroquinone (DQ), an electron scavenger, for its ability to sequester excess electrons during OGD. Low-dose DQ provided moderate neuroprotection and reduced mitochondrial DNA release (Fig. 2J-2L); however, higher concentrations induced substantial neuronal death. Thus, its narrow therapeutic window limits its potential as a stroke treatment. Together, these findings establish electron overload as a central driver of reductive stress during acute cerebral ischemia and highlight the need for safe strategies to remove excess electrons at their source.

To clarify how electron accumulation alters enzymatic reactions through NAD(H) imbalance, we performed metabolic flux analysis in neurons supplemented with NADH. An elevated NADH/NAD□ ratio increased nearly all glycolytic metabolites (fig. S3A-S3E). While serine synthesis, a glycolytic branch pathway requiring NAD□ as a cofactor, was not enhanced by NADH treatment (Fig. S3F). In contrast, TCA cycle metabolites exhibited distinct responses: most intermediates decreased, whereas the entry substrates acetyl-CoA and citrate were preserved (fig. S3G-S3N). Notably, succinate accumulated drastically after NADH exposure (fig. S3L). These findings demonstrate that electron overload broadly rewires cellular metabolism by disrupting NAD(H) homeostasis. Consistently, NADH treatment upregulated key metabolic enzymes, including HK2, PC, and SDH (fig. S3R), highlighting reciprocal regulation between metabolic flux and redox state. Because OGD induces reductive stress in neurons, we next examined its metabolic consequences. OGD suppressed core glycolysis (fig. S3S) and downstream branching pathways, including the pentose phosphate pathway (fig. S3T) and serine synthesis pathway (fig. S3U). Citrate and α-ketoglutarate, two major TCA intermediates, were also markedly reduced (fig. S3V). In contrast, succinate levels remained unchanged after OGD in neuron (fig. S3V), further analysis of the downstream conversion of TCA cycle intermediates revealed robust accumulation of succinate (fig. S3W). These results indicate both substrate depletion and limited availability of electron acceptors.

### Fructose enters neurons via GLUT5 and KHK-catalyzed reactions

Ischemic preconditioning (IPC) is an effective strategy against ischemic stroke (*27, 28*). To explore how IPC alleviates reductive stress during ischemia, we performed RNA-seq and LC/MS analyses on mouse brain tissues and detected prominent alterations in metabolic pathways (Figs. 3A-3C). Notably, the level of fructose-1-phosphate (F1P), a key metabolite in fructose metabolism, was markedly elevated in the injured brain region of IPC-treated mice (Fig. 3C). In MCAO mice, fructose abundance increased substantially in the lesion area at 1 h post-ischemia, and the mRNA level of Aldob, a core enzyme for fructose metabolism, was upregulated following respiratory chain suppression (fig. S4A). A previous study also reported higher fructose levels in hypoxia-tolerant naked mole-rats relative to mice (*13*). These findings led us to hypothesize that fructose metabolism is enhanced under hypoxic conditions. To test this, we measured the expression of enzymes involved in fructose and mannose metabolism in MCAO mouse brains and OGD-exposed neurons, and observed significant upregulation of these genes (fig. S8B). Collectively, these data indicate that fructose metabolism is selectively activated in response to ischemia-induced mitochondrial dysfunction.

**Fig. 3.**
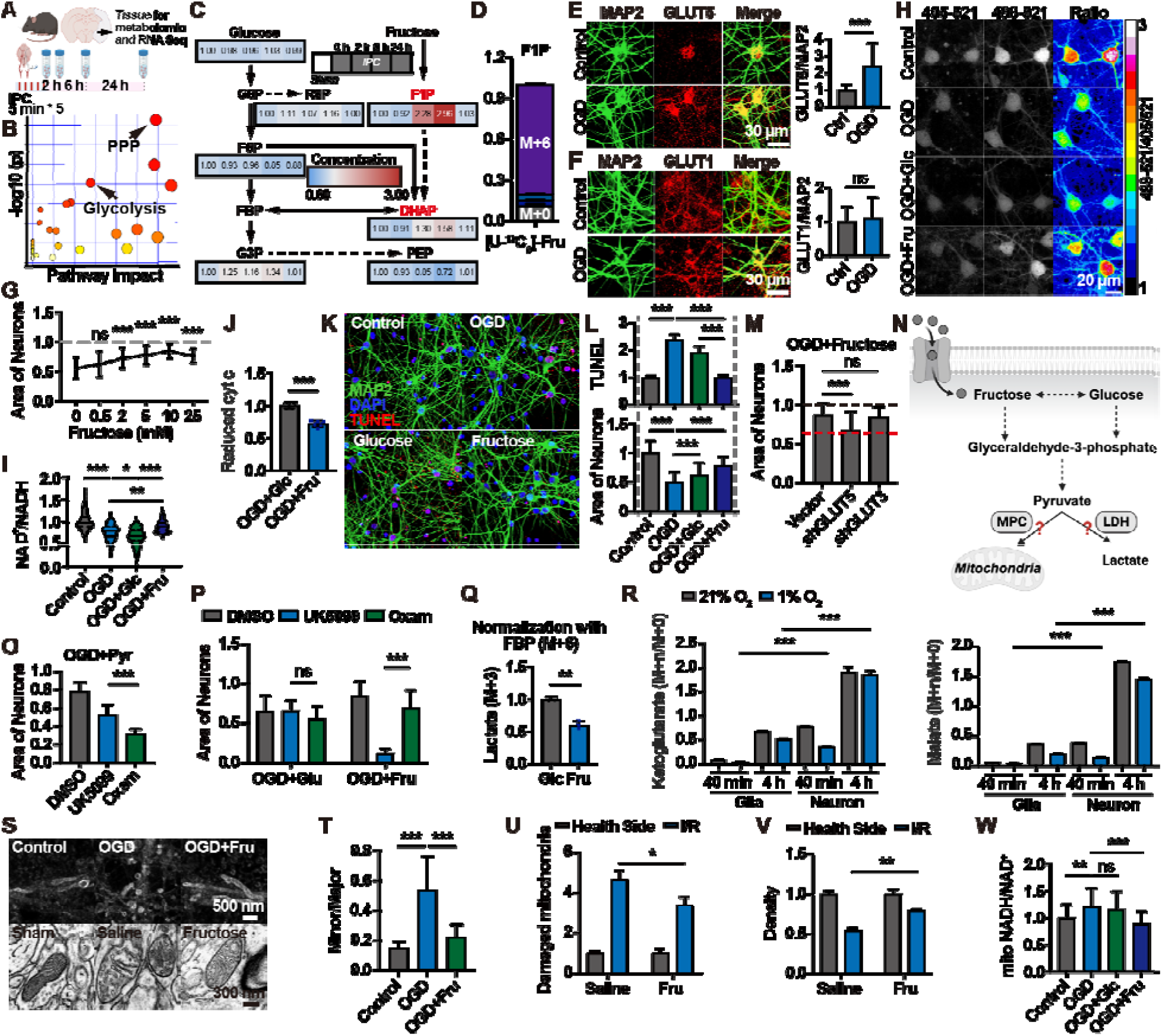
Fructose enters neurons via GLUT5 and exerts mitochondrial-protective effect reliant on mitochondrial metabolic turnover. **(A-C)** Mice received five cycles of 5-min ischemic preconditioning. Brain tissue was harvested at 0 h, 2 h, 6 h, and 24 h post-preconditioning for subsequent RNA-seq (B) and metabolomic profiling (C). **(D)** Primary neurons were incubated with 10 mM U-¹³C fructose for 4 h, and downstream fructose-1-phosphate (F1P) abundance was quantified via LC-MS (N=4). **(E and F)** Primary neurons were exposed to 4 h OGD to recapitulate ischemic injury. Immunofluorescent co-staining of GLUT5 (E) or GLUT1 (F) with the neuronal marker MAP2 allowed quantitative measurement of protein abundance (N=4, n>6). **(G)** Primary neurons were exposed to OGD-reperfusion with a gradient of fructos concentrations supplemented during OGD; MAP2 immunostaining was used to evaluate neuronal morphological integrity (N=4, n>5). **(H and I)** Primary neurons expressing the SoNar redox biosensor were were exposed to 4 h OGD, SoNar fluorescence signals were quantified (N=5, n>6). **(J)** Primary neurons were exposed to 4 h OGD with 10 mM glucose or 10 mM fructose provided. Reduced cytochrome c content was measured immediately upon OGD termination (N=6). **(K and L)** Primary neurons were exposed to 4 h OGD with 10 mM glucose or 10 mM fructose provided, followed by 24 h reperfusion. TUNEL, MAP2, and DAPI co-staining was applied to label apoptotic neurons; the proportion of TUNEL /MAP2 /DAPI cells were quantified to compare apoptotic rates across groups (N=4, n>6). **(M)** Primary neurons were separately transfected with GLUT5-targeting or GLUT3-targeting shRNAs. Cell viability was quantified after 4 h OGD exposure, and values were normalized to normoxic control neurons expressing shGLUT5 or shGLUT3 (N=4, n>5). **(N)** Schematic illustrating divergent fates of glycolytic pyruvate: mitochondrial oxidative metabolism versus cytosolic lactate fermentation. **(O)** During OGD-reperfusion, neurons were supplemented with pyruvate together with DMSO, UK5099, or oxamate; all compounds were removed upon reperfusion, and neuronal viability was statistically assessed (N=6, n>5). **(P)** During OGD-reperfusion, neurons were supplemented with glucose or fructose together with DMSO, UK5099, or oxamate; all compounds were removed upon reperfusion, and neuronal viability was statistically assessed (N=4, n>5). **(Q)** Primary neurons were treated with 10 mM U-¹³C glucose or U-¹³C fructose for the full 4 h OGD period. LC-MS was used to measure M+3 lactate levels, normalized to M+6 fructose-1,6-bisphosphate (FBP) abundance (N=3). **(R)** Primary astrocytes and neurons were incubated with U-¹³C□ glucose tracer under normoxic (21% O□) or hypoxic (1% O□) conditions for either 40 min or 4 h. LC-MS quantification determined the fractional ratios of labeled versus unlabeled α-ketoglutarate and malate pools (N=4). **(S-U)** Stimulated emission depletion (STED) super-resolution microscopy of cultured neurons and transmission electron microscopy (TEM) of mouse brain tissue were performed to characterize fructose-mediated alterations to mitochondrial ultrastructure (neuron: N=4, n>10; brain tissue: N=3, n>3). **(V)** Transmission electron microscopy was performed on brain tissues from C57BL/6 mice subjected to tMCAO surgery to quantify damaged mitochondria and cellular structural density. **(W)** Primary neurons transduced with mitochondrially targeted MTS-SoNar were subjected to 4 h OGD in glucose- or fructose-supplemented medium. Intramitochondrial NADH/NAD□ ratios were quantified at the 4 h OGD endpoint for each treatment group (N=6, n>5). Data are presented as means ± SD (E, F, G, I, J, L, M, O, P, Q, R, T, W) or SEM (U and V). Multiple-group comparisons were performed using one-way analysis of variance (ANOVA), followed by Dunnett’s post-hoc test (G, M, O, T and W) or Tukey’s HSD post-hoc test (I and L); two-way analysis of variance (ANOVA) was performed, followed by Sidak’s multiple comparisons test (P, R, U and V); comparisons between two groups were conducted using the unpaired Student’s t-test (E, F, J and Q). *P < 0.05, **P < 0.01, ***P < 0.001; ns, not significant.

Stable isotope tracing demonstrated that exogenous fructose could enter cultured neurons and is converted to F1P (Fig. 3D), although the underlying transport mechanism remains unclear. Several glucose transporter family members, including GLUT2, GLUT5, GLUT7, GLUT9, and GLUT11, mediate fructose uptake, with GLUT5 displaying the highest substrate specificity (*29*). We observed that GLUT5 was selectively upregulated in neurons following OGD both in vitro and in vivo (Figs. 3E, 3F, fig. S4C-S4G). Since ischemic neurons suffer from both hypoxia and metabolic substrate deprivation (*30–32*), supplemented fructose may protect injured neurons as an ischemic metabolic substrate. Indeed, we found fructose supplementation protected OGD-injured neurons in a dose-dependent manner, with 10 mM displaying maximal protection (Fig. 3G). Unlike glucose, fructose markedly reduced the NADH/NAD□ ratio, attenuated mitochondrial reduced cytochrome c accumulation (Fig. 3H-3J), and suppressed OGD-induced neuronal death as assessed by TUNEL and MAP2 staining (Figs. 3K, 3L). These findings indicate that, although both sugars serve as metabolic substrates, they exert distinct effects on redox homeostasis.

To determine whether fructose uptake is required for neuron protection, we inhibited GLUT5 using its competitive antagonist 2,5-anhydro-D-mannitol (2,5-AM) and found that 2,5-AM markedly abolished fructose-mediated neuroprotection (fig. S4H) and reduced the incorporation of ¹³C -fructose into downstream metabolites (fig. S4J-SJM). In contrast, inhibition of GLUT1 exerted little effect on neuronal viability (fig. S4I). To further validate the neuroprotective effects of fructose, we knocked down GLUT5 and GLUT3 in neurons (fig. S4N-S4P). GLUT5 knockdown did not alter neuronal viability under normal culture conditions. GLUT3 silencing reduced cell survival under standard culture conditions (fig. S4Q, S4R), since GLUT3 acts as the primary glucose transporter in neurons (*33, 34*). In OGD-injured neurons, GLUT5 knockdown largely abrogated fructose-mediated neuroprotection, whereas GLUT3 depletion had no impact on this protective effect (all data were normalized to normoxic GLUT5 and GLUT3 knockdown neurons, respectively) (Fig. 3M). Collectively, fructose enters neurons via GLUT5 during OGD, and its distinct metabolic profile restores neuronal NAD(H) homeostasis, conferring neuroprotection.

Metabolomic profiling revealed that fructose fuels rapid and robust production of downstream glycolytic intermediates relative to glucose, with metabolite pools plateauing at the 4-hour time point (fig. S5B, S5D-J). Fructose can be converted to glucose and then phosphorylated to glucose-6-phosphate by hexokinase 2 (HK2) for glycolysis or metabolized to fructose-1-phosphate (F1P) by ketohexokinase (KHK) (*35–38*) (fig. S4R), a hallmark that distinguishes fructose metabolism from glucose metabolism. We found that the HK2 inhibitor 2-deoxy-D-glucose (2-DG) barely affected fructose-treated neurons, whereas KHK inhibition markedly abolished fructose-mediated protection (fig. S4S). Furthermore, we found that fructose treatment further elevated Aldob levels in neurons subjected to OGD. Inhibition of glucose 6 phosphate dehydrogenase (G6PD) by 6 aminonicotinamide (6 AN) exerted minimal effects on neuronal viability (fig. S4T), and exogenous NADPH failed to protect neurons (fig. S4U, S4V). Fructose also did not alter the NADPH/NADP ratio (fig. S4W) or increase ribulose-5-phosphate (R5P) levels (fig. S5C). Collectively, the PPP is not the primary pathway mediating fructose-induced neuroprotection. Likewise, neither serine nor glycine supplementation improved neuronal viability (fig. S4X). Inhibition of serine hydroxymethyltransferase 1/2 (SHMT1/2) with SHIN1 exerted no obvious effects on cell viability (fig. S4Y), and rescue with serine or glycine did not alter survival after serine synthesis blockade (fig. S4Z). Moreover, fructose treatment for 4 h did not increase cellular serine content (fig. S5D). These results indicate that the serine synthesis pathway is not required for fructose-mediated neuroprotection.

### Fructose metabolism relies on mitochondria and alleviates mitochondrial damage

Pyruvate is either fermented to lactate or transported into mitochondria for catabolism (Fig. 3N). Lactate fermentation regenerates NAD□, whereas mitochondrial TCA cycle metabolism consumes NAD□ and produces NADH. To determine how distinct pyruvate fates affect neuronal survival under OGD, we treated OGD-exposed neurons with pyruvate together with DMSO, UK5099, or oxamate. Inhibition of lactate fermentation by oxamate produced more severe neuronal injury (Fig. 3O). We found that fructose treatment yielded a lower NADH/NAD□ ratio than glucose treatment (Fig. 3H, 3I), suggesting that fructose might alleviate reductive stress and confer neuroprotection by promoting lactate fermentation. To test this, we applied DMSO, UK5099, or oxamate to fructose- or glucose-treated neurons. Unexpectedly, UK5099 markedly reduced the viability of fructose-supplemented neurons (Fig. 3P), and isotope tracing analyses revealed that fructose produced far less lactate than glucose (Fig. 3Q, S5E), indicating that fructose preferentially undergoes mitochondrial metabolism.

This result contradicted our initial hypothesis. To examine mitochondrial metabolic activity under hypoxia, we exposed neurons and glial cells (as a control) to 21% or 1% oxygen for 40 min or 4 h in the presence of U-¹³C□-glucose, followed by LC/MS-based metabolite quantification. After 4 h of hypoxia, neurons exhibited substantially higher isotopic labeling of mitochondrial metabolites, including α-ketoglutarate and malate, compared with glial cells (Fig. 3R, figs. S6H-S6J, S6L, S6M), whereas glycolytic metabolites showed no such difference (fig. S6B, S6D, S6E). Consistently, ETC blockade with antimycin A failed to further induce transcriptional upregulation of core glycolytic genes in primary neurons (fig. S6O). Together, these data demonstrate that neurons sustain active mitochondrial metabolism even under hypoxia, while lactate fermentation persists.

To further investigate the impacts of fructose on mitochondria, we evaluated neuronal mitochondrial morphology. Fructose alleviated OGD-induced mitochondrial injury both in vitro and in vivo (Fig. 3S-3U, Fig. S7J) and attenuated the disruption of cellular structural integrity (Fig. 3V). Furthermore, fructose inhibited mitochondrial DNA leakage and cardiolipin depletion, maintained inner mitochondrial membrane fluidity (fig. S7D, S7E), and blunted the rise in mitochondrial permeability (fig. S7G), likely by lowering the mitochondrial NADH/NAD□ ratio relative to glucose treatment (Fig. 3W). Interestingly, glucose supplementation increased mitochondrial matrix proton accumulation during OGD (fig. S7H), which arises from the generation of NADH and protons during glucose metabolism and may perturb mitochondrial membrane potential. In contrast, proton accumulation was negligible after fructose treatment, indicating distinct metabolic identity between the two sugars. Sodium ions also contribute to mitochondrial membrane potential (*39*), yet matrix sodium levels remained unaltered in neurons under OGD with or without glucose or fructose supplementation (fig. S7I). Given that the TCA cycle generates large amounts of NADH along with proton. Thus, fructose and glucose may differentially modulate the TCA cycle, leading to distinct effects on intramitochondrial proton abundance.

### Fructose sustains NAD(H) homeostasis by driving reverse TCA cycle flux

After transport into mitochondria, Pyruvate enters the TCA cycle via two routes: conversion to acetyl-CoA by pyruvate dehydrogenase (PDH), or generation of oxaloacetate by pyruvate carboxylase (PC) (Fig. 4A). To determine which pathway mediates fructose-induced neuroprotection, we inhibited PC or PDH in fructose-treated neurons. Only ZY-444 (PC inhibitor) markedly reduced neuronal survival (Fig. 4B), indicating that PC-dependent fructose metabolism sustains TCA cycle activity and redox homeostasis (*40*). Notably, exogenous oxaloacetate failed to rescue PC-deficient neurons, possibly because oxaloacetate cannot freely cross the inner mitochondrial membrane (*41, 42*). The forward TCA cycle, with PDH catalyzing its initial step, consumes NAD□ to generate NADH (*43, 44*) (*45*), whereas the reverse TCA cycle, whose first reaction is mediated by PC, exerts the opposite effect (*46–48*). We sought to determine whether the protective effect of fructose in blunting the OGD-induced elevation of the NADH/NAD□ ratio (Fig. 3W, fig. S7K-S7M) is dependent on TCA cycle flux direction. To this end, we pharmacologically inhibited malate dehydrogenase (MDH) using a specific MDH inhibitor (LW6). MDH suppression reduces NADH production during forward TCA cycling but lowers NAD□ generation when the cycle runs in reverse (Fig. 4A). We found that MDH inhibition did not change the NAD□/NADH ratio in glucose-treated neurons but markedly decreased this ratio in fructose-supplemented cells (Fig.4D, 4E). These results demonstrate that fructose maintains NAD(H) homeostasis by promoting reversal of the TCA cycle.

**Fig. 4.**
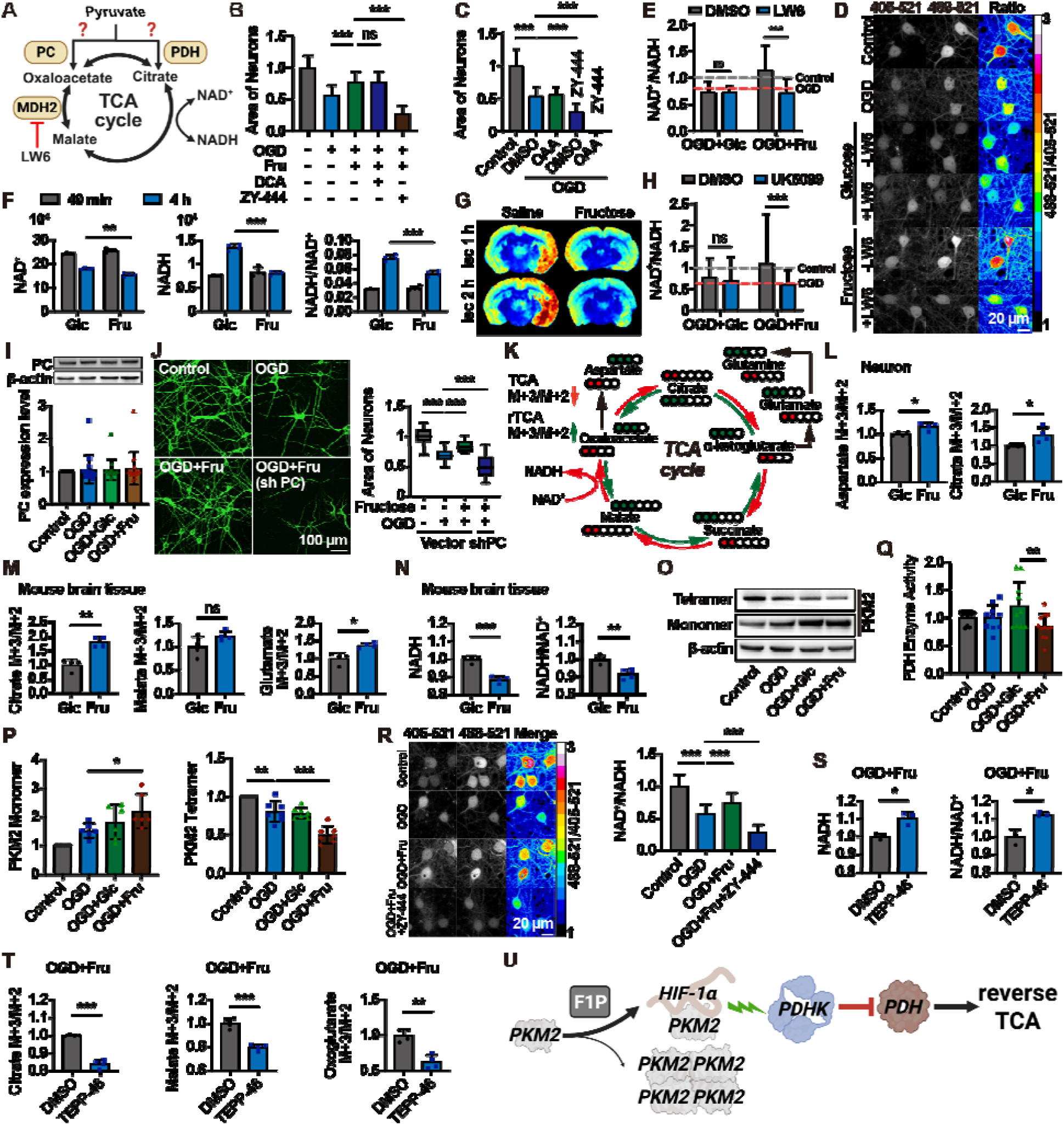
Fructose influx mitigates reductive stress under oxygen-glucose deprivation by stimulating reductive carboxylation through TCA cycle anaplerosis. **(A)** Schematic illustrating two fates of glycolytic pyruvate: anaplerotic conversion to oxaloacetate catalyzed by pyruvate carboxylase (PC), or entry into the TCA cycle as acetyl-CoA via pyruvate dehydrogenase (PDH). Malate dehydrogenase (MDH) serves as a core TCA-cycle enzyme; pharmacological MDH blockade disrupts cellular NAD(H) homeostasis by impairing TCA cycle turnover. **(B)** Quantification of neuronal survival using MAP2 immunofluorescence. Primary neurons received 4 h OGD co-supplemented with vehicle DMSO, 10 mM dichloroacetate (DCA), or 5 µM ZY-444. All compounds were washed out, and cultures underwent a 24 h reoxygenation recovery period prior to viability assessment (N=6, n>6). **(C)** Primary neurons were exposed to OGD-reperfusion injury. During OGD, cultures were treated with 2 mM oxaloacetate (OAA) alongside DMSO or 15 µM ZY-444; MAP2 immunostaining was performed to calculate neuronal survival rates across groups (N=3, n>5). **(D and E)** Ratiometric quantification of the intracellular NAD□/NADH ratio via the SoNar biosensor in primary cortical neurons. After 4 h OGD with vehicle DMSO or malate dehydrogenase inhibitor (MADi), live-cell real-time imaging was conducted to profile dynamic NAD(H) redox status (N=6, n>8). **(F)** Primary neurons were incubated with glucose or fructose throughout 4 h OGD. LC-MS metabolomic measurement was performed immediately post-OGD to quantify absolute NAD□ and NADH abundance as well as the resulting NADH/NAD□ ratio (N=4). **(G)** Mice subjected to tMCAO received intravenous saline or fructose at ischemia onset. Brain tissue was harvested at 1 h and 2 h post-ischemia for mass spectrometry imaging to map spatial fluctuations in endogenous NADH abundance within the injured lesion. **(H)** Primary cortical neurons expressing the SoNar redox biosensor underwent 4 h OGD, followed by metabolic rescue with glucose or fructose in the presence of DMSO or 10 µM UK5099. The intracellular NAD□/NADH ratio was quantified via ratiometric SoNar fluorescence readouts (N=7, n>10). **(I)** Primary cortical neurons were exposed to 4 h OGD, and whole-cell lysates were harvested for western blot (WB) analysis of pyruvate carboxylase (PC) protein expression (N=15). **(J)** Primary neurons with lentivirus-mediated shRNA knockdown of PC (shPC) were exposed to OGD with 10 mM fructose supplementation. MAP2 immunofluorescence staining was used to quantify neuronal viability (N=4, n>5). **(K)** Stochastic computational modeling of mass isotopomer distributions derived from ¹³C-labeled glucose or fructose enables quantitative dissection of forward versus reductive TCA cycle flux. **(L)** Primary neurons were cultured with [U-¹³C□]-glucose or [U-¹³C□]-fructose during 4 h OGD. LC-MS analysis was implemented to calculate M+3/M+2 isotopic fractional ratios for aspartate and citrate (N=4). **(M)** At the onset of tMCAO occlusion, mice received an intravenous bolus of 93 mg/kg [U-¹³C□]-glucose or [U-¹³C□]-fructose. One hour later, the injured ipsilateral cerebral hemisphere was dissected for LC-MS quantification of M+3/M+2 ratios for key TCA-cycle intermediates: citrate, malate, and glutamate (N=4). **(N)** Mice undergoing tMCAO were administered 90 mg/kg glucose or fructose intravenously upon ischemia initiation. Ipsilateral lesioned brain tissue was collected at 1 h post-occlusion for LC-MS measurement of total NADH levels and the NADH/NAD□ ratio. **(O and P)** Primary neurons were exposed to 4 h OGD with glucose or fructose supplementation. Western blot assays were conducted to compare the abundance of monomeric and tetrameric PKM2 protein across experimental groups (tetrameric PKM2 protein: N=7, monomeric PKM2 protein: N=6 (1 outlier excluded from initial n=7)). **(Q)** Primary neurons received 10 mM glucose or 10 mM fructose throughout 4 h OGD. Total cellular PDH enzymatic activity was measured via commercial colorimetric assay kits immediately after OGD termination (N=5, n>=2). **(R)** Primary neurons were transduced with the SoNar biosensor to report intracellular NAD□/NADH ratios. Cultures were subjected to 4 h OGD with fructose co-treatment plus either DMSO or ZY-444, and endpoint SoNar ratiometric fluorescence signals were recorded (N=4, n>5). **(S and T)** Fructose was supplemented to neurons during OGD rescue, alongside vehicle DMSO or TEPP-46. LC-MS profiling was performed to quantify NADH abundance, NADH/NAD□ ratios, and M+3/M+2 isotopic fractions of core TCA-cycle metabolites in each treatment cohort (N=4). **(U)** Fructose catabolism generates fructose-1-phosphate (F1P), which locks PKM2 into its monomeric conformation. This monomeric PKM2 shift rewires mitochondrial metabolic networks and drives reverse flux through mitochondrial PC. Data are presented as means ± SD. Multiple-group comparisons were performed using one-way analysis of variance (ANOVA), followed by Dunnett’s post-hoc test (B, C, I, J, P and R) or Tukey’s HSD post-hoc test (Q); two-way analysis of variance (ANOVA) was performed, followed by Sidak’s multiple comparisons test (E, F and H); comparisons between two groups were conducted using the unpaired Student’s t-test (L, M, N, S and T). *P < 0.05, **P < 0.01, ***P < 0.001; ns, not significant.

To further validate fructose mediated regulation of NAD(H) homeostasis, we performed LC/MS analysis in OGD neurons treated with fructose or glucose. Fructose reduced NADH levels and the NADH/NAD□ ratio (Fig. 4F). Furthermore, via mass spectrometry imaging of brain tissue, we verified that fructose alleviates NADH accumulation within the injured cerebral regions in MCAO mice (Fig. 4G). Using SoNar probe, we found that inhibition of mitochondrial fructose metabolism with UK5099 abolished this regulatory effect (Fig. 4H), indicating that fructose maintains redox homeostasis primarily via mitochondria. Furthermore, we found that although PC expression remained comparable between glucose-and fructose-treated neurons under OGD (Fig. 4I), PC knockdown in neurons (fig. S7N) compromised cell viability (fig.S7O, S7P). After normalization to the viability of non-OGD neurons, PC silencing largely abrogated fructose-mediated protection (Fig. 4J). Exogenous NAD□ partially rescued the survival of PC-deficient neurons subjected to OGD (fig. S7Q), suggesting the PC pathway modulates NAD(H) homeostasis. Collectively, fructose relieves reductive stress under OGD, likely by entering the TCA cycle in a PC-dependent manner to sustain NAD(H) homeostasis, with minimal impacts on GSH and GSSG levels (fig. S7R).

### F1P functions as a metabolic signal to activate non-canonical reverse TCA cycling

To directly determine TCA cycle direction during fructose metabolism, we performed isotope tracing in OGD neurons using [U-¹³C□] -glucose and [U-¹³C□]-fructose. Forward TCA cycling yields metabolites such as citrate, oxaloacetate and glutamate predominantly with two labeled carbons (M+2)(*47, 49*), whereas reverse TCA flux produces metabolites carrying three labeled carbons (M+3) (Fig. 4K). Fructose treatment markedly increased the M+3/M+2 ratio of TCA intermediates both in vitro and in vivo (Fig. 4L, 4M). Consistently, fructose reduced NADH levels and the NADH/NAD□ ratio in the ischemic brain lesion compared with glucose (Fig. 4N). Collectively, fructose alleviates reductive stress by promoting reverse TCA cycle flux.

We next explored how fructose drives reverse TCA cycling. Previous studies have demonstrated that fructose-1-phosphate (F1P) inhibits tetramerization of pyruvate kinase M2 (PKM2) (*50, 51*). PKM2 monomers suppress pyruvate dehydrogenase (PDH) activity (*52*), rerouting pyruvate toward pyruvate carboxylase (PC)-dependent TCA metabolism. We thus hypothesized that F1P derived from fructose acts as a signaling molecule to trigger TCA cycle reversal and mitigate reductive stress. We found that fructose significantly increased PKM2 monomer abundance while reducing PKM2 tetramer levels (Fig. 4O, 4P), and concurrently inhibited PDH activity (Fig. 4Q). These results indicate that fructose maintains PKM2 in its monomeric state and suppresses forward TCA cycle flux. Moreover, PC blockade abolished the ability of fructose to lower the NADH/NAD□ ratio (Fig. 4R), confirming that fructose fuels the reverse TCA cycle in a PC-dependent manner. To verify the role of PKM2 monomers in fructose-driven reverse TCA cycling, OGD neurons with fructose treatment were incubated with DMSO or TEPP-46, a compound that stabilizes PKM2 tetramers. TEPP-46 markedly attenuated fructose-induced TCA cycle reversal (Fig. 4T) and elevated NADH content as well as the NADH/NAD□ ratio (Fig. 4S). In summary, fructose is catabolized to F1P, which acts as a signaling molecule to inhibit PDH activity. This metabolic rewiring diverts pyruvate to the PC-dependent reverse TCA cycle (Fig. 4U). By serving as an energy substrate and restoring NAD(H) homeostasis via this pathway, fructose exerts potent neuroprotective effects.

### Fructose exerts prominent cerebral protective effects in mice

Given the complex architecture of the brain, including the blood-brain barrier (BBB) and tripartite synapses, we first examined whether fructose could reach injured brain regions in the tMCAO mouse model. We intraperitoneally injected [U-¹³C□]-glucose or [U-¹³C□]-fructose immediately after ischemia onset. Brain tissues were collected for mass spectrometry imaging 1-hour post-ischemia. Isotope-labeled downstream metabolites were detected in the lesioned area, confirming that fructose can be metabolized in damaged brain tissue (fig. S5L). Encouraged by the potent neuroprotective effects of fructose in OGD models, we next assessed its therapeutic potential in vivo. TTC staining of brain sections from tMCAO mice revealed that fructose treatment markedly reduced cerebral infarct volume (Fig. 5A, 5B). This protective effect was consistently observed in both aged and diabetic mice (Fig. 5C, 5D). By contrast, glucose or galactose exerted no neuroprotection, while sucrose conferred a mild protective effect (fig. S8A, S8B). This mild benefit is likely attributed to fructose produced from sucrose hydrolysis. Since TTC staining relies on succinate dehydrogenase (SDH) to convert colorless TTC into red formazan, fructose, as a glycolytic substrate, might alter SDH activity and interfere with the accuracy of infarct assessment. To exclude this possibility, we performed magnetic resonance imaging (MRI) in tMCAO mice, which confirmed that fructose significantly reduced brain lesion area (Fig. 5E) and cerebral edema (fig. S8J). Furthermore, we found that fructose increased the number of MAP2-positive surviving neurons within the injured region (Fig. 5F), validating its potent neuroprotective effect. In addition, Fructose significantly improved neurological deficit scores following ischemia-reperfusion injury (fig. S8H). Electroencephalography (EEG) recordings showed that fructose barely affected EEG energy but markedly increased wave amplitude (fig. S8K-S8M). Epileptiform discharges and overt seizures were detected in saline-treated mice during the ischemic phase (fig. S8N, S8O), whereas these abnormal electrical activities were absent in fructose-treated animals. These findings indicate that fructose preserves normal neuronal electrical activity. In line with our findings, PC inhibition largely abolished the cerebral protection conferred by fructose (Fig. 5G), demonstrating that the in vivo protective action of fructose is PC-dependent.

**Fig. 5.**
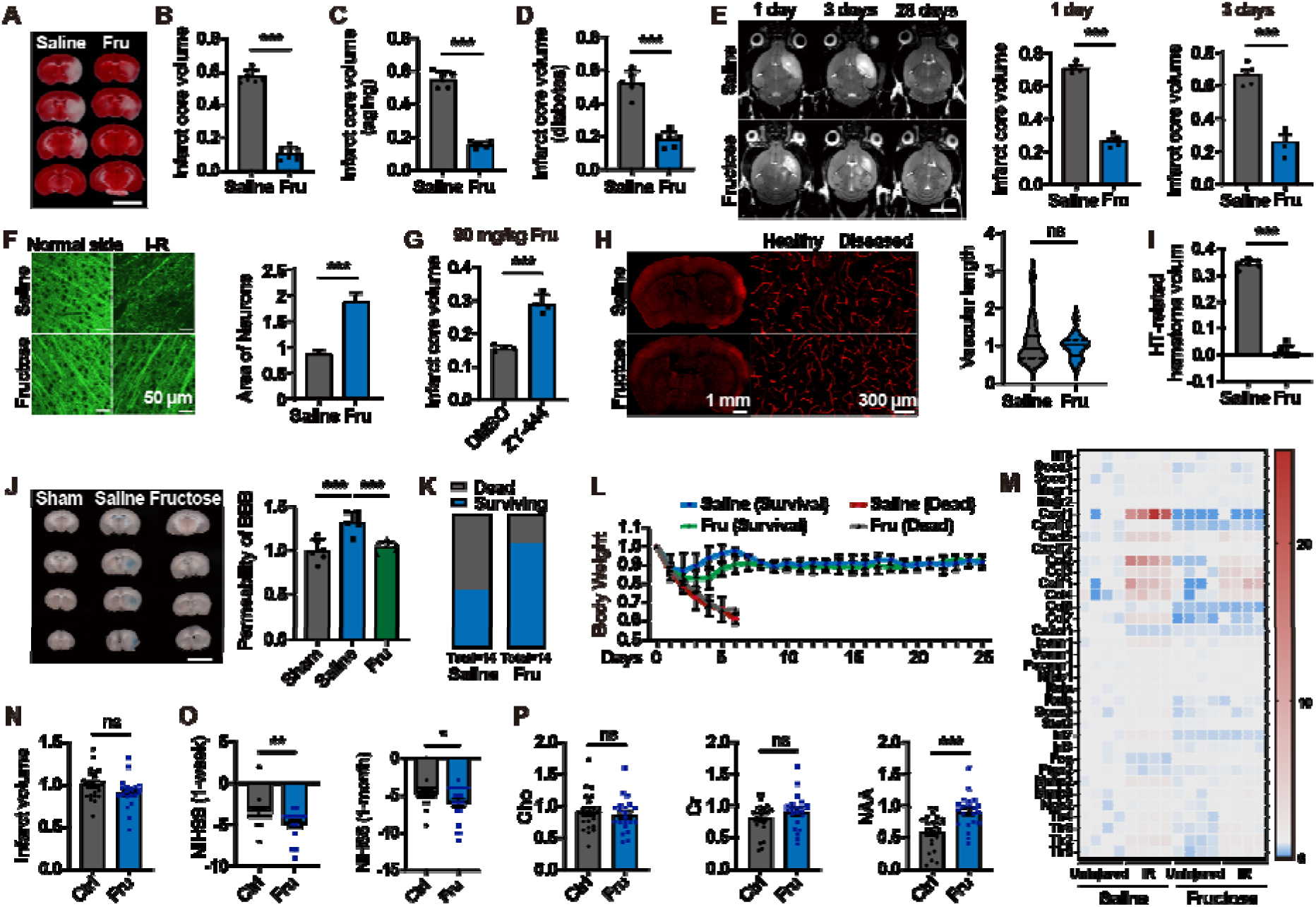
Fructose intervention elicits robust neuroprotective effects against cerebral ischemia-reperfusion injury in mice. **(A-D)** Twelve-week-old wild-type male C57BL/6 mice (N=8) (A, B), aged 12-month-old wild-type male C57BL/6 mice (N=6) (C), and 12-week-old male BKS-Leprem2Cd479/Gpt mice (N=6) (D) were subjected to transient middle cerebral artery occlusion (tMCAO) and treated with vehicle saline or 90 mg/kg fructose. Cerebral infarct volume across all cohorts was quantified via 2,3,5-triphenyltetrazolium chloride (TTC) staining (scale bar=5 mm). **(E)** Serial 9.4 T multi-parametric T2-weighted MRI scans were acquired at 1, 3, and 28 days after tMCAO in two experimental groups: vehicle-treated ischemic controls and mice receiving 90 mg/kg fructose intervention (N>=4). **(F)** MAP2 immunofluorescence staining was used to quantify surviving neurons within the cortical infarct territory of tMCAO-operated mice (N=5, n>3). **(G)** TTC staining at 24 h post-tMCAO assessed changes in cerebral infarct volume in mice administered fructose alone or fructose co-delivered with ZY-444 (N=4). **(H)** Vascular remodeling was visualized 72 h after reperfusion in saline- and fructose-treated tMCAO mice via systemic perfusion with FITC-conjugated Lycopersicon esculentum lectin to label cerebral microvessels (N=6, n>3). **(I)** Mice subjected to tMCAO received intravenous saline or 90 mg/kg fructose at ischemia onset; brain tissue was harvested at 24 h post-reperfusion to evaluate hemorrhagic transformation (N=6). **(J)** Evans Blue (EB) extravasation assays were performed at 24 h post-reperfusion to quantitatively evaluate blood-brain barrier (BBB) permeability in tMCAO mice (N=6) (scale bar=5 mm). **(K)** Survival analysis revealed 28-day post-ischemia survival rates in untreated tMCAO mice versus in cohorts receiving 90 mg/kg fructose (N=14). **(L)** Longitudinal tracking of body weight changes in fructose-treated tMCAO mice relative to saline vehicle controls (N>6). **(M)** RNA-seq profiling uncovered divergent expression patterns of neuroinflammation-associated genes within peri-infarct cerebral tissue at 1-month post-tMCAO (N=4). **(N)** Diffusion-weighted imaging (DWI) was applied to human stroke patients receiving standard care alone or standard care combined with fructose adjunct therapy. Infarct volume was quantified by comparing gray-scale signal intensity between lesioned brain regions and matched contralateral mirror regions (Standard thrombolysis: 23 patients, Standard thrombolysis plus fructose treatment: 21 patients). **(O)** Quantification of relative reductions in National Institutes of Health Stroke Scale (NIHSS) scores at 1 week and 1-month post-treatment versus baseline values in control patients and patients receiving fructose intervention (Standard thrombolysis: 23 patients, Standard thrombolysis plus fructose treatment: 21 patients). **(P)** In vivo magnetic resonance spectroscopy (MRS) was used to quantify relative shifts in cerebral choline (Cho), creatine (Cr), and N-acetylaspartate (NAA) within damaged brain tissue relative to uninjured contralateral regions in control and fructose-treated human subjects (Standard thrombolysis: 23 patients, Standard thrombolysis plus fructose treatment: 21 patients). For mouse experiments, data are presented as means ± SD. Multiple-group comparisons were performed using one-way analysis of variance (ANOVA), followed by Dunnett’s post-hoc test (J); Comparisons between two groups were conducted using the unpaired Student’s t-test (B, C, D, E, F, G, H and I); For clinical analyses, data are presented as means ± SEM. Between-group comparisons were conducted via Mann-Whitney U test (N, O and P). *P < 0.05, **P < 0.01, ***P < 0.001; ns, not significant.

To determine whether fructose mediates cerebral protection through alternative pathways, we sequentially assessed cerebral perfusion, inflammatory responses, and ATP levels in mouse brain tissue. Using laser speckle imaging at multiple time points in tMCAO mice, we found that fructose exerted no detectable influence on cerebral perfusion (fig. S8C). Vascular labeling with lectin (Fig. 5H) and CD31 staining (fig. S8I) further confirmed that fructose exerted minimal effects on cerebral vessels. However, fructose markedly reduced hemorrhagic transformation and blood-brain barrier leakage (Fig. 5I, 5J). Survival statistics revealed that fructose significantly reduced the mortality rate from cerebral ischemia-reperfusion injury (Fig. 5K), while no notable change in body weight was observed (Fig. 5L). By examining inflammatory cell infiltration in mouse brain tissues at 1 and 7 days after tMCAO, we observed that, relative to saline treatment, fructose did not alter the total number of microglia or pro-inflammatory microglia (fig. S8D-S8G). RNA-seq analysis of brain samples collected one month after ischemia-reperfusion further demonstrated that fructose did not upregulate the expression of inflammation-related genes (Fig. 5M). Collectively, fructose treatment does not exacerbate inflammatory responses. By performing in vivo two-photon imaging in tMCAO mice, we found that neuronal ATP gradually declined during ischemia and recovered upon reperfusion (fig. S8P). LC/MS and mass spectrometry imaging confirmed reduced ATP and ADP contents during ischemia (fig. S8Q, S8S), while the ATP/ADP ratio rose markedly at 2 hours post-ischemia (fig. S8R). Phosphate creatine levels decreased only at 1 hour of ischemia but increased substantially at 2 hours, and total creatine remained unchanged throughout ischemia (fig. S8Q). Fructose treatment exerted no significant effects on the levels of these metabolites (fig. S8S, S8T).

### Fructose provides neuroprotection for patients with acute ischemic stroke

With the safety and neuroprotective efficacy of fructose treatment validated in primary neuronal cultures and multiple mouse models (including wild-type, diabetic, and aged mice animals), fructose therapy may improve clinical outcomes. We conducted a single-center, prospective, randomized, open-label, blinded-endpoint (PROBE) phase IIa exploratory proof-of-concept trial (ClinicalTrials.gov identifier NCT07626541). A total of 53 patients were screened for eligibility, and 44 participants eligible for intravenous thrombolysis were included in the intention-to-treat (ITT) analysis: 23 patients received standard thrombolysis alone (control group), while 21 patients received 10% fructose infusion initiated prior to thrombolysis alongside standard reperfusion therapy (intervention group; Table S1, Fig. S9). Safety surveillance focused on intracranial hemorrhage, hepatorenal function, glycemic fluctuation and neurological deterioration over the 30-day follow-up. The incidence of adverse events was comparable between groups (Table S2), with no unanticipated treatment-related serious adverse events identified. The relative change in NIHSS score at 7-day post-thrombolysis was the prespecified primary endpoint. Predefined secondary exploratory outcomes included serial 7-day and 30-day NIHSS, 30-day modified Rankin Scale (mRS), 24 h post-recanalization DWI infarct volume and edema, and cerebral metabolites measured by ¹H-MRS.

Assessment of DWI images acquired 24 hours after recanalization showed no significant reduction in infarct severity with fructose treatment (P= 0.0740) (Fig. 5N), which may be attributed to the limited sample size and substantial individual heterogeneity. Nevertheless, fructose markedly reduced NIHSS scores at 1 week (P= 0.0016) and 1 month (P= 0.0099) after reperfusion (Fig. 5O, fig. S8U), with a correlation between fructose treatment and sustained NIHSS improvement (fig. S8X). No significant intergroup differences were observed for 30-day mRS distribution or favorable recovery rates (Fig. S8V, S8W, Table S3). Divergent findings between the NIHSS and mRS may reflect inherent characteristics of each scale: the continuous NIHSS captures evolving acute neurological deficits, whereas the ordinal mRS is influenced by multiple confounding factors, including rehabilitation training, spontaneous brain plasticity, complications, and underlying comorbidities, which may mask group differences. Additionally, the mRS represents a broad categorical scale for global disability and is less sensitive to mild functional improvements than the NIHSS. Collectively, although no statistically significant difference was found in long-term activities of daily living, the remarkable improvement in acute neurological function still carries important clinical implications. Proton MRS of ischemic tissue demonstrated comparable choline (involved in the synthesis and breakdown of cell membrane phospholipids) and creatine levels across groups (Fig. 5P), consistent with preclinical mouse findings (Fig. S8T). By contrast, N-acetylaspartate (NAA), a biomarker of neuronal integrity mainly synthesized within neuronal mitochondria, showed higher concentrations in the fructose-treated group (P = 0.0064), indicative of attenuated ischemic neuronal injury. Collectively, these exploratory clinical findings yield preliminary human evidence supporting a potential neuroprotective effect of adjunctive fructose therapy among stroke patients undergoing thrombolysis. Larger confirmatory trials are needed to validate these findings prior to consideration of clinical implementation.

## DISCUSSION

Many existing stroke studies center on antioxidant strategies designed to eliminate reperfusion-derived ROS. However, conventional antioxidants such as edaravone show limited therapeutic efficacy (*53, 54*), raising an important question: do neurons suffer additional forms of injury before the ROS surge? Our findings identify ischemia-induced reductive stress within the infarct region as an early driver of neuronal injury. During oxygen deprivation, excessive accumulation of reducing equivalents impairs NADH oxidation and NAD□ regeneration, disrupting neuronal redox homeostasis and metabolic activity. Such reductive imbalance causes cardiolipin depletion, mitochondrial inner membrane destabilization, and mtDNA release, thereby promoting neuronal injury.

To counteract hypoxia-induced reductive stress, cells employ multiple adaptive mechanisms, including NAD□ regeneration through lactate fermentation (*55*), utilization of alternative electron acceptors such as fumarate (*24*), producing 2-oxoglutarate (2OG) (*56*), and redistribution of reducing equivalents through mitochondrial shuttle systems such as the reverse malate-aspartate shuttle (*57*) and SLC25A51 (*58*). Beyond these endogenous responses, targeted mitochondrial rewiring may offer other therapeutic opportunities to restore redox balance. We identify reverse TCA cycle activation as an effective strategy to resolve ischemia-associated reductive stress. Notably, fructose metabolism generates fructose-1-phosphate (F1P), which redirects pyruvate into a pyruvate carboxylase–dependent reverse TCA cycle, thereby promoting NADH oxidation while replenishing TCA intermediates to sustain ischemic metabolism. Although the tissue sources and biochemical origins of ischemic precondition-induced fructose production remain unclear, these findings suggest that fructose metabolism may represent an evolutionarily conserved metabolic adaptation that enhances hypoxia tolerance (*13*).

Given that NAD(H) homeostasis can be modulated by inhibiting NAD□degradation (*59, 60*), enhancing NAD□biosynthesis (*61, 62*), or supplementing exogenous NAD□, alternative strategies to counteract reductive stress may exist. However, whether these approaches can rapidly correct the pathological NADH/NAD□imbalance during ischemia remains unknown. Moreover, exogenous NAD□supplementation can further increase NADH levels (*19*), requires mitochondrial transport, and may have limited metabolic impact when functional substrates are depleted during ischemia, thereby restricting its neuroprotective potential. In contrast, fructose may provide two distinct advantages: it directly restores NADH/NAD□ balance by remodeling TCA cycle flux and re-establishing electron transfer, while also providing metabolic substrate to sustain neuronal metabolism during ischemia. We demonstrate that intravenously administered fructose could efficiently reaches the brain and neurons, and protects against ischemic injury in wild-type, aged, and diabetic mice. In our exploratory clinical trial, adjunctive fructose treatment improved neurological recovery, reduced NIHSS scores, and restored cerebral metabolite profiles associated with neuronal integrity. Although imaging-based outcomes did not reach statistical significance, likely due to limited sample size and interindividual variability, larger cohorts may reveal additional structural benefits. Together, our findings establish reductive stress as an upstream metabolic vulnerability during ischemia and suggest that restoring redox flux presents an exciting area for future research.

## Acknowledgments

We thank members of the Zhou lab for critical discussion and technical assistance; Wenzhi Su and Woo-ping Ge, Chinese Institute for Brain Research, Beijing (CIBR), for discussion fructose cell specificity; We thank Professor Qionghai Dai and Jiamin Wu, Tsinghua University, for the help of with DAOSLMIT-2P imaging framework; We are also grateful to Zhongshuang Lv and Xixia Li for helping with sample preparation and taking TEM images at the Center for Biological Imaging (CBI), Institute of Biophysics, Chinese Academy of Science; Fengting Jiang and Qi Tang for the MALDI-TOF Mass Spectrometry image at Guidon Pharmaceutics Ltd.; We also thank Yupei Jiao and YusongWang at Facility center of Metabolomics and lipidomics in Tsinghua University for their help with the LC-MS/MS experiments; We also thank BioRender (https://BioRender.com) for the academic publication license and editable scientific drawing templates, which facilitated the construction of schematic mechanism figures presented in this work.

## Author contributions

T.-C.R. designed and executed the preclinica experiments, analyzed the data, and wrote the original manuscript; Feng Yu designed the clinical study, performed patient enrollment, and drafted the clinical-related sections of the manuscript; J.L. performed ischemic-precondition-omic study; Y.-X.S performed the virus injection and 2P imaging on mice; Y-L.W helped interpret the results; and X.-M.J. co-designed and initiated the project; B.Z. is the senior author conceived the supervised the project. T.-C.R. and B.Z. wrote the manuscript with the help of the other authors.

## Supplementary Materials

Materials and Methods

Supplementary Text

Figs. S1 to S9

Tables S1 to S3

## References and Notes

1. M. Bélanger, I. Allaman, P. J. Magistretti, Brain energy metabolism: focus on astrocyte-neuron metabolic cooperation. Cell Metab 14, 724–738 (2011).

2. D. Vilchez et al., Mechanism suppressing glycogen synthesis in neurons and its demise in progressive myoclonus epilepsy. Nat Neurosci 10, 1407–1413 (2007).

3. M. Erecińska, I. A. Silver, ATP and brain function. J Cereb Blood Flow Metab 9, 2–19 (1989).

4. T. Kalogeris, C. P. Baines, M. Krenz, R. J. Korthuis, Cell biology of ischemia/reperfusion injury. Int Rev Cell Mol Biol 298, 229–317 (2012).

5. E. T. Chouchani et al., Ischaemic accumulation of succinate controls reperfusion injury through mitochondrial ROS. Nature 515, 431–435 (2014).

6. K. Zhang et al., Hollow Prussian Blue Nanozymes Drive Neuroprotection against Ischemic Stroke via Attenuating Oxidative Stress, Counteracting Inflammation, and Suppressing Cell Apoptosis. Nano letters 19, 2812–2823 (2019).

7. Y. Tian et al., Silencing of TXNIP Alleviated Oxidative Stress Injury by Regulating MAPK-Nrf2 Axis in Ischemic Stroke. Neurochemical research 45, 428–436 (2020).

8. S. Lorenzano et al., Early molecular oxidative stress biomarkers of ischemic penumbra in acute stroke. Neurology 93, e1288–e1298 (2019).

9. D. Zhang et al., 9-Methylfascaplysin exerts anti-ischemic stroke neuroprotective effects via the inhibition of neuroinflammation and oxidative stress in rats. International immunopharmacology 97, 107656 (2021).

10. Y. N. Kim et al., Neuroprotective effects of PEP-1-carbonyl reductase 1 against oxidative-stress-induced ischemic neuronal cell damage. Free Radic Biol Med 69, 181–196 (2014).

11. R. Rodrigo et al., Oxidative stress and pathophysiology of ischemic stroke: novel therapeutic opportunities. CNS Neurol Disord Drug Targets 12, 698–714 (2013).

12. J. J. Lochhead, P. T. Ronaldson, T. P. Davis, The role of oxidative stress in blood-brain barrier disruption during ischemic stroke: Antioxidants in clinical trials. Biochem Pharmacol 228, 116186 (2024).

13. T. J. Park et al., Fructose-driven glycolysis supports anoxia resistance in the naked mole-rat. Science (New York, N.Y.) 356, 307–311 (2017).

14. Y. Zhao et al., SoNar, a Highly Responsive NAD+/NADH Sensor, Allows High-Throughput Metabolic Screening of Anti-tumor Agents. Cell Metab 21, 777–789 (2015).

15. S. Norcross et al., Extending roGFP Emission via Förster-Type Resonance Energy Transfer Relay Enables Simultaneous Dual Compartment Ratiometric Redox Imaging in Live Cells. ACS Sens 2, 1721–1729 (2017).

16. W. Xiao, J. Loscalzo, Metabolic Responses to Reductive Stress. Antioxid Redox Signal 32, 1330–1347 (2020).

17. W. Hummel, B. Riebel, Isolation and biochemical characterization of a new NADH oxidase from Lactobacillus brevis. Biotechnol Lett 25, 51–54 (2003).

18. D. V. Titov et al., Complementation of mitochondrial electron transport chain by manipulation of the NAD+/NADH ratio. Science 352, 231–235 (2016).

19. R. P. Goodman et al., Hepatic NADH reductive stress underlies common variation in metabolic traits. Nature 583, 122–126 (2020).

20. X. Pan et al., A genetically encoded tool to increase cellular NADH/NAD(+) ratio in living cells. Nat Chem Biol 20, 594–604 (2024).

21. W. Li et al., Release of damaged mitochondrial DNA: A novel factor in stimulating inflammatory response. Pathol Res Pract 258, 155330 (2024).

22. J. Trnka, F. H. Blaikie, R. A. Smith, M. P. Murphy, A mitochondria-targeted nitroxide is reduced to its hydroxylamine by ubiquinol in mitochondria. Free Radic Biol Med 44, 1406–1419 (2008).

23. C. Li et al., Mito-Tempo prevents nicotine-induced exacerbation of ischemic brain damage. J Appl Physiol (1985) 125, 49–57 (2018).

24. J. B. Spinelli et al., Fumarate is a terminal electron acceptor in the mammalian electron transport chain. Science 374, 1227–1237 (2021).

25. W. X. Ma, C. Y. Li, R. Tao, X. P. Wang, L. J. Yan, Reductive Stress-Induced Mitochondrial Dysfunction and Cardiomyopathy. Oxid Med Cell Longev 2020, 5136957 (2020).

26. M. R. de Oliveira, Dimethyl fumarate and mitochondrial physiology: implications for neurological disorders. Front Pharmacol 17, 1748360 (2026).

27. C. Hou et al., Chronic remote ischaemic conditioning in patients with symptomatic intracranial atherosclerotic stenosis (the RICA trial): a multicentre, randomised, double-blind sham-controlled trial in China. Lancet Neurol 21, 1089–1098 (2022).

28. W. Zhao et al., Safety and Efficacy of Remote Ischemic Preconditioning in Patients With Severe Carotid Artery Stenosis Before Carotid Artery Stenting: A Proof-of-Concept, Randomized Controlled Trial. Circulation 135, 1325–1335 (2017).

29. S. A. Oppelt, W. Zhang, D. R. Tolan, Specific regions of the brain are capable of fructose metabolism. Brain Res 1657, 312–322 (2017).

30. C. E. Murry, R. B. Jennings, K. A. Reimer, Preconditioning with ischemia: a delay of lethal cell injury in ischemic myocardium. Circulation 74, 1124–1136 (1986).

31. J. M. Gidday, Cerebral preconditioning and ischaemic tolerance. Nature reviews. Neuroscience 7, 437–448 (2006).

32. R. Meng et al., Upper limb ischemic preconditioning prevents recurrent stroke in intracranial arterial stenosis. Neurology 79, 1853–1861 (2012).

33. S. Nagamatsu et al., Neuron-specific glucose transporter (NSGT): CNS distribution of GLUT3 rat glucose transporter (RGT3) in rat central neurons. FEBS letters 334, 289–295 (1993).

34. W. Peng et al., Glucose transporter 3 in neuronal glucose metabolism: Health and diseases. Metabolism 123, 154869 (2021).

35. G. Marek et al., Adiponectin resistance and proinflammatory changes in the visceral adipose tissue induced by fructose consumption via ketohexokinase-dependent pathway. Diabetes 64, 508–518 (2015).

36. T. Hayasaki et al., Fructose increases the activity of sodium hydrogen exchanger in renal proximal tubules that is dependent on ketohexokinase. The Journal of nutritional biochemistry 71, 54–62 (2019).

37. C. Patel et al., Effect of dietary fructose on portal and systemic serum fructose levels in rats and in KHK-/- and GLUT5-/- mice. American journal of physiology. Gastrointestinal and liver physiology 309, G779–790 (2015).

38. G. T. Cori, S. Ochoa, M. W. Slein, C. F. Cori, The metabolism of fructose in liver; isolation of fructose-I-phosphate and inorganic pyrophosphate. Biochimica et biophysica acta 7, 304–317 (1951).

39. P. Hernansanz-Agustín et al., A transmitochondrial sodium gradient controls membrane potential in mammalian mitochondria. Cell 187, 6599–6613.e6521 (2024).

40. D. A. Cappel et al., Pyruvate-Carboxylase-Mediated Anaplerosis Promotes Antioxidant Capacity by Sustaining TCA Cycle and Redox Metabolism in Liver. Cell metabolism 29, 1291–1305.e1298 (2019).

41. I. G. Darvey, Does the transport of oxaloacetate across the inner mitochondrial membrane during gluconeogenesis require carrier proteins other than those used in the malate-aspartate shuttle? Biochemical education 28, 80–82 (2000).

42. S. Passarella, F. Palmieri, E. Quagliariello, The transport of oxaloacetate in isolated mitochondria. Archives of biochemistry and biophysics 180, 160–168 (1977).

43. A. Moussaieff et al., Glycolysis-mediated changes in acetyl-CoA and histone acetylation control the early differentiation of embryonic stem cells. Cell metabolism 21, 392–402 (2015).

44. Z. Cai et al., Phosphorylation of PDHA by AMPK Drives TCA Cycle to Promote Cancer Metastasis. Molecular cell 80, 263–278.e267 (2020).

45. G. Tournaire et al., Skeletal progenitors preserve proliferation and self-renewal upon inhibition of mitochondrial respiration by rerouting the TCA cycle. Cell reports 40, 111105 (2022).

46. M. J. Lu et al., SLC25A51 decouples the mitochondrial NAD(+)/NADH ratio to control proliferation of AML cells. Cell metabolism 36, 808–821.e806 (2024).

47. A. R. Mullen et al., Reductive carboxylation supports growth in tumour cells with defective mitochondria. Nature 481, 385–388 (2011).

48. C. Sun et al., NAD depletion mediates cytotoxicity in human neurons with autophagy deficiency. Cell reports 42, 112372 (2023).

49. A. Xie et al., Citrate clearance is a major function of aconitase 2 in the canonical TCA cycle. Cell 189, 2684–2699.e2621 (2026).

50. S. R. Taylor et al., Dietary fructose improves intestinal cell survival and nutrient absorption. Nature 597, 263–267 (2021).

51. W. Luo et al., Pyruvate kinase M2 is a PHD3-stimulated coactivator for hypoxia-inducible factor 1. Cell 145, 732–744 (2011).

52. T. Golias et al., Hypoxic repression of pyruvate dehydrogenase activity is necessary for metabolic reprogramming and growth of model tumours. Scientific reports 6, 31146 (2016).

53. P. A. Lapchak, A critical assessment of edaravone acute ischemic stroke efficacy trials: is edaravone an effective neuroprotective therapy? Expert Opin Pharmacother 11, 1753–1763 (2010).

54. S. Feng et al., Edaravone for acute ischaemic stroke. Cochrane Database Syst Rev 2011, Cd007230 (2011).

55. H. C. D. Medeiros, S. Y. Lunt, The Warburg effect: Saturation of mitochondrial NADH shuttles triggers aerobic lactate fermentation. Mol Cell 82, 3119–3121 (2022).

56. W. M. Oldham, C. B. Clish, Y. Yang, J. Loscalzo, Hypoxia-Mediated Increases in L-2-hydroxyglutarate Coordinate the Metabolic Response to Reductive Stress. Cell Metab 22, 291–303 (2015).

57. P. Altea-Manzano et al., Reversal of mitochondrial malate dehydrogenase 2 enables anaplerosis via redox rescue in respiration-deficient cells. Mol Cell 82, 4537–4547.e4537 (2022).

58. T. S. Luongo et al., SLC25A51 is a mammalian mitochondrial NAD(+) transporter. Nature 588, 174–179 (2020).

59. J. Camacho-Pereira et al., CD38 Dictates Age-Related NAD Decline and Mitochondrial Dysfunction through an SIRT3-Dependent Mechanism. Cell Metab 23, 1127–1139 (2016).

60. H. Y. McGuinness, W. Gu, Y. Shi, B. Kobe, T. Ve, SARM1-Dependent Axon Degeneration: Nucleotide Signaling, Neurodegenerative Disorders, Toxicity, and Therapeutic Opportunities. Neuroscientist 30, 473–492 (2024).

61. S. Srivastava, Emerging therapeutic roles for NAD(+) metabolism in mitochondrial and age-related disorders. Clin Transl Med 5, 25 (2016).

62. S. Imai, A possibility of nutriceuticals as an anti-aging intervention: activation of sirtuins by promoting mammalian NAD biosynthesis. Pharmacol Res 62, 42–47 (2010).

