## Supplementary Material for "Reductive stress contributes to neuronal injury in ischemic stroke: fructose protection via the reverse TCA cycle"

**The PDF file includes:**

Materials and Methods

Figs. S1 to S9

Tables S1 to S3

**Materials and Methods**

Primary Neuron Culture

Cortical primary neuronal cultures were generated from postnatal day 0 (P0) SD rat pups harvested within 12 h of delivery. Animals were sacrificed via cold-induced narcosis for 2-3 min before whole-brain dissection. Meningeal membranes were fully stripped, cortical tissues were dissected into small fragments, and enzymatic dissociation was performed using papain incubation at 37 °C for 45 min. Following centrifugation, the supernatant was discarded, and the cell pellet was resuspended in inhibitor solution. The cell suspension was then mixed with 1% bovine serum albumin (BSA) solution and centrifuged again. After removal of the supernatant, cells were resuspended in complete culture medium consisting of Neurobasal-A Medium, B27 Supplement, GlutaMAX Supplement, fetal bovine serum (FBS), and penicillin-streptomycin. Finally, neurons were seeded at desired densities and incubated in a CO₂ incubator.

Primary Astrocyte Culture

Primary astrocytes were isolated from postnatal day 2 (P2) Sprague-Dawley (SD) rat pups. Animals were subjected to cold narcosis on ice for 3-5 min until fully unconscious, after which the whole brain tissues were rapidly dissected. Following complete removal of meningeal membranes, hippocampal tissues were excised and minced into small fragments, then enzymatically digested in trypsin solution at 37 °C for 12 min. After centrifugation, the supernatant was aspirated, and tissue pellets were resuspended in culture medium containing high-glucose DMEM supplemented with 10% fetal bovine serum (FBS) and 1% penicillin-streptomycin (PS). The tissue suspension was mechanically dissociated via repeated pipetting to eliminate intact tissue aggregates. Purified astrocytes were seeded at required densities and maintained in a humidified CO₂ incubator.

293T Cell Culture

HEK 293T cells were cultured in high-glucose DMEM supplemented with 10% fetal bovine serum (FBS) and 1% penicillin-streptomycin (PS). Cells were maintained at 37 °C in a humidified incubator with 5% CO₂. For virus packaging, culture dishes were pre-coated with Matrigel before cell seeding. Virus packaging was performed when cells reached approximately 80% confluence.

Lentivirus production and Lentivirus infection

The target gene sequence was inserted into the lentiviral transfer plasmid pFUGW for subsequent virus packaging. HEK293T cells were co-transfected with three plasmids (transfer vector pFUGW, packaging plasmids psPAX2 and pMD2.G) at a 5:3:2 mass ratio using Lipofectamine 3000 transfection reagent in accordance with the manufacturer’s protocols. Cell culture medium was refreshed 20 h post-transfection to eliminate residual transfection complexes. Culture supernatant containing lentiviral particles was harvested at 48 h after medium replacement, filtered through a 0.45 μm sterile filter to remove cell debris, and aliquoted for long-term storage at -80 °C. Freshly isolated neurons (6 × 10⁵ cells) were incubated with 1 mL of the lentiviral supernatant.

shRNA-mediated gene knockdown

Lentiviral short hairpin RNA (shRNA) constructs targeting the indicated gene (*sh-PC-1* CACCTATGGTTTCCCTATTATCTCGAGATAATAGGGAAACCATAGGTGTTTTTT; *sh-PC-2* CAAGCCCATCAAGAAAGTAATCTCGAGATTACTTTCTTGATGGGCTTGTTTTTT; *sh-PC-3* ACTACTTCATCGAGGTCAATTCTCGAGAATTGACCTCGATGAAGTAGTTTTTTT; *sh-Slc2a5* CCGGTGGTTGGATTCTTGGTGAATACTCGAGTATTCACCAAGAATCCAACCATTTTTT; *sh-Slc2a3* CCGGGCAGGACTTTCGAGGATATTACTCGAGTAATATCCTCGAAAGTCCTGCTTTTTT) and a non-targeting scramble shRNA control were used for gene knockdown experiments. For in vitro neuronal knockdown, primary neurons were transfected with lentiviral particles. Subsequent functional experiments were performed after 7 days of knockdown to achieve sufficient gene silencing efficiency. Western blotting were routinely applied to verify the knockdown efficiency at both protein and mRNA levels.

Hypoxia Treatment and Hypoxia signal Measurement

Neuronal cultures were transferred to a STEMCELL hypoxic incubation chamber (Cat. 27311). A premixed gas blend (1% O₂, 5% CO₂, 94% N₂) was flushed through the system at a constant 25 L/min flow rate for 5 min, after which all inlet and outlet valves were tightly closed, and sealed incubation proceeded at 37 °C.

Cell immunofluorescence

Cellular samples underwent fixation in 4% paraformaldehyde (PFA) for 15 min, followed by three sequential PBS washes. Non-specific antigen binding was blocked with blocking buffer for 1 h before overnight primary antibody incubation at 4 °C. Samples were washed three times in PBS, then exposed to fluorophore-conjugated secondary antibodies for 1 h at ambient temperature under light-shielded conditions. Samples were rinsed with PBS and double-distilled water (ddH₂O), mounted, and imaged using a confocal laser scanning microscope. For subsequent TUNEL staining, samples were washed three times with PBS after incubation with secondary antibodies, and then incubated with TUNEL working solution (Beyotime, Cat. No. C1090) for 1 h. After PBS rinsing, samples were mounted with DAPI-containing mounting medium and examined under a confocal microscope.

Measurement of NAD^+^/NADH

Cells were plated in 96-well plates. NAD^+^/NADH was measured using a modified version of the manufacturer’s instructions supplied with the NAD/NADH Glo Assay (Promega).

Measurement of PDH Enzyme Activity

Cells were plated in 3.5 cm dishes. PDH Enzyme Activity was measured using a modified version of the manufacturer’s instructions supplied with the Pyruvate dehydrogenase (PDH) Enzyme Activity Microplate Assay Kit (abcam).

Western blot analysis

Cellular pellets were lysed on ice for 30 min in NP-40 lysis buffer supplemented with protease and phosphatase inhibitor cocktails. Lysates were clarified by 13,000 g centrifugation (5 min), and soluble protein fractions were retained. Total protein quantification was performed using the BCA assay, and the samples were heated in SDS-PAGE loading buffer at 100 °C for 10 min to denature the proteins. An equal amount of protein was loaded in sodium dodecyl sulfate-polyacrylamide gel electrophoresis (SDS-PAGE) and transferred to a polyvinylidene fluoride (PVDF) membrane. The membrane was blocked in 5% BSA and incubated with primary antibodies at 4℃ overnight with mild agitation. After incubation with secondary antibodies at room temperature for 1 h, the signal intensities were visualized and analyzed using ImageJ.

Tissue immunofluorescence

Tissue sections were fixed with 4% paraformaldehyde for 20 min, followed by three rinses with phosphate-buffered saline (PBS). After 1 h of blocking, the sections were incubated with primary antibodies overnight at 4 °C. After three washes in PBS, secondary antibodies were applied and incubated for 1 h at room temperature in the dark. The sections were then rinsed with PBS, mounted, and visualized using a confocal laser scanning microscope.

Reduced Cytochrome c Assay

Eight-day-cultured neurons were subjected to oxygen-glucose deprivation (OGD) for 4 h. Upon completion of OGD treatment, cells were harvested under continuous hypoxia. The cells were dissociated with TrypLE Express (Gibco, 2444890) in a hypoxic environment and transferred to centrifuge tubes. Samples were centrifuged at 300 rcf for 3 min, and the cell pellets were resuspended in OGD Buffer under hypoxic conditions. The cell suspension was then loaded into a quartz cuvette. To isolate the system from ambient air, mineral oil was overlaid on the buffer solution. Characteristic absorbance peaks at 550 nm and 530 nm were measured using a UV spectrophotometer, and all data were normalized to the absorbance at 530 nm.

Neuronal Metabolic Flux and Metabolomics Sample Preparation

For metabolic flux analysis, [U-¹³C₆]-glucose or [U-¹³C₆]-fructose was dissolved in glucose-free OGD buffer to prepare a 10 mM working solution. The original neuronal culture medium was then replaced with the isotope-labeled OGD buffer. For metabolomic profiling, neuronal culture medium was exchanged with OGD buffer supplemented with unlabeled 10 mM glucose or fructose. Subsequently, the cells were placed in a hypoxic chamber and incubated at 37 °C for 4 h in the dark. After incubation, samples were immediately quenched with liquid nitrogen. A cold extraction solution consisting of acetonitrile:methanol:water (4:4:2, v/v/v) was added to the samples. Cells were scraped off on ice and transferred together with the extraction solution into centrifuge tubes. The mixtures were kept at −80 °C for no less than 2 h, followed by centrifugation at 12,000 rcf for 10 min at 4 °C. The resulting supernatants were collected into pre-chilled tubes for subsequent LC/MS analysis.

LC/MS Analysis of Metabolites Involved in Glycolysis and TCA Cycle

Isotope labeling data acquisition was performed using a Shimadzu LC system coupled to a Triple TOF mass spectrometer (QTOF 6600+, AB Sciex, made in Woodlands Central Industrial Estate, Singapore). Chromatographic separation was achieved using a HILIC column (IHILIC-(P) Classic column, 5 μm, 150mm × 2.1mm, 200 A, made in Sweden). The mobile phase A was 20 mmol/L ammonium acetate, 0.1% ammonium hydroxide, and 2.5 μmol/L methylene diphosphate in 95:5 water:ACN, and mobile phase B was ACN. The gradient was as follows: 0min, 85% B; 2min, 85% B; 7min, 65% B; 12min, 35% B; 12.1 min, 20% B; 15.9 min, 20% B; 16 min, 85% B; 23min, 85% B. The flow rate was set at 0.2mL/min with a sam-ple injection volume of 5 μL and the total run time at 23min. ESI parameters setup was GS1, 60; GS2, 60; CUR, 35; temperature, 500; ISVF, −4500 in negative modes. For each metabolite, the areas of 12C and various 13C isotopologue peaks were integrated using El-MAVEN software (version 0.12.1). Correction for natural isotope abundance was performed using R package AccuCor. The labeling fraction of a specific isotopologue was calculated as the area of that isotopo-logue divided by the sum of all isotopologues’ area.

Fructose-1-phosphate Detection

The Dionex Ultimate 3000 UPLC system was coupled to a TSQ Quantiva Ultra triple-quadrupole mass spectrometer (Thermo Fisher, CA), equipped with a heated electrospray ionization (HESI) probe. Extracts were separated by a synergi Hydro-RP column (2.0×100mm, 2.5 μm, phenomenex). A binary solvent system was used, in which mobile phase A consisted of 10 mM tributylamine adjusted with 15 mM acetic acid in water, and mobile phase B of methanol. This analysis used a 25-minute gradient from 5% to 90% mobile B. Data acquired in selected reaction monitoring (SRM) for metabolites in positive-negative ion switching mode. The resolution for Q1 and Q3 are both 0.7 FWHM. The source voltage was 3500v for positive and 2500v for negative ion mode. The source parameters are as follows: capillary temperature: 350^o^C; heater temperature: 300^o^C; sheath gas flow rate: 35; auxiliary gas flow rate: 10. Tracefinder 3.2 (Thermo, USA) was applied for metabolite identification and peak integration. Mass spectrometry standard of fructose-1-phosphate was used for qualitative and quantitative detection of metabolites in the experiment.

Mitochondrial Permeability Assay

Neurons were pre-transfected with lenti-mito DsRed to label mitochondria. Neurons subjected to different treatments were incubated in OGD buffer containing 1 μM Calcein-AM for 15 min in the dark. After removal of the staining solution, cells were rinsed twice with OGD buffer. The cells were then incubated in OGD buffer supplemented with 1 mM CoCl₂ for 30 min under dark conditions. Following two additional washes with OGD buffer, images were acquired using a confocal microscope, and the fluorescence intensity of intramitochondrial Calcein-AM was quantified.

Mitochondrial Isolation

Mitochondria were isolated from primary cultured neurons. Briefly, a total of 1 × 10⁷ neurons were dissociated with TyrpLETM Express (Gibco, 2444890) and homogenized using a glass tissue grinder. Mitochondrial purification was subsequently performed following the manufacturer’s protocols of the commercial kit (Cat. No. PK10016). The isolated mitochondria were used for subsequent measurements of matrix protons and sodium ions.

Intramitochondrial Proton and Sodium Ion Measurement

This probe labeling protocol was modified according to a previously published method (*39*). Isolated mitochondrial fractions were incubated at 37 °C for 20 min with 10 μM 2',7'-bis-(2-carboxyethyl)-5-(and-6)-carboxyfluorescein acetoxymethyl ester (BCECF-AM) or 1,3-benzenedicarboxylic acid, 4,4'-[1,4,10-trioxa-7,13-diazacyclopentadecane-7,12-benzofurandiyl)]bis-, tetrakis [(acetyloxy)methyl] ester (SBFI-AM). BCECF-AM was used to detect matrix hydrogen ions, while SBFI-AM was applied for matrix sodium ion measurement. Unbound dyes were removed by two rounds of centrifugation at 12,000 g for 5 min at 4 °C, and the mitochondrial pellets were resuspended in sucrose buffer. Mitochondria labeled with BCECF-AM were loaded into 24-well glass-bottom plates fitted with No. 1.5 high-profile coverslips (thickness: 0.17 ± 0.005 mm; Cat. No. P24-1.5H-N). The plates were centrifuged at 7000 g for 15 min at 4 °C to attach mitochondria to the bottom of each well. Fluorescence signals of BCECF-AM were acquired using an Andor Dragonfly 200 confocal microscope (Model: CR-DFLY-202-2540). The excitation wavelengths were set at 390 nm and 485 nm, with the emission wavelength fixed at 530 nm for both channels. Mitochondria stained with SBFI-AM were aliquoted into 96-well plates containing sodium-free buffer and measured using a CLARIOstar multi-mode microplate reader. For SBFI-AM detection, excitation wavelengths were 355 nm and 390 nm, and the emission wavelength was set to 530 nm.

Super-resolution Imaging of Mitochondria

Neurons from each experimental group were incubated with OGD buffer containing 200 μM PK Mito Red at 37 °C for 15 min in the dark. After incubation, the staining solution was removed, and cells were rinsed twice with OGD buffer. Cells were briefly fixed with 2.5% glutaraldehyde for 30 s, followed by further fixation with 4% paraformaldehyde (PFA) for 15 min. Upon completion of fixation, cells were washed three times with PBS and imaged using a STED super-resolution microscope.

Mitochondrial Membrane Fluidity Assay

Neurons treated with different conditions were incubated in OGD Buffer containing 200 nM CMTMRos for 15 minutes in the dark. The dye solution was then aspirated, and cells were rinsed twice with OGD Buffer. Subsequently, the mitochondrial membrane structure was subjected to fluorescence photobleaching under a confocal microscope. Time-lapse imaging was acquired immediately after photobleaching, and the fluorescence recovery rate of the bleached region was calculated.

RNA Extraction

For cellular samples, 2×10⁶ neurons were seeded into 6 cm culture dishes. After oxygen-glucose deprivation (OGD) treatment, total RNA was extracted using the RNAprep Pure Cultured Cell/Bacteria Total RNA Extraction Kit (Cat. No. DP430). For tissue samples, mice were deeply anesthetized and perfused. Brain tissues were rapidly harvested and stored at −80 °C. During RNA extraction, 20 mg of tissue was mixed with 300 μL of lysis buffer and fully homogenized using a tissue homogenizer. Subsequent procedures were performed according to the manufacturer’s instructions of the RNAprep Pure Tissue Kit (Cat. No. DP431).

Animal

The BeiHang University Biomedical Ethics Committee in Beijing, China, approved all animal studies. All experimental procedures were performed in accordance with the National Institutes of Health guidelines for the Care and Use of Laboratory Animals. The C57BL/6J mice and SD rats were purchased from SiPeiFu, and the BKS-Leprem2Cd479/Gpt mice were purchased from Vital River. Mice were housed in a vivarium under a 12-h light/dark cycle at 22℃, with unrestricted access to food and water. Animals were randomly assigned to each treatment group, and all experiments were conducted in accordance with standard laboratory procedures for randomization and blinding. Mouse euthanasia was performed by carbon dioxide (CO_2_) inhalation followed by cervical dislocation.

Transient middle cerebral artery occlusion model

Twelve-week-old male C57BL/6 mice weighing 25-27 g were enrolled for transient MCAO surgeries; monofilament sutures were size-matched to individual animal weight. Following inhalation anesthesia, mice were secured in a supine position on a temperature-controlled heating pad that maintained core temperature at 37 °C, and a midline cervical vertical cut was created using microdissection scissors. The common carotid artery, internal carotid artery, and external carotid artery were isolated. The monofilament suture was inserted into the internal carotid arterya (At this time point, mice were intraperitoneally injected with 90 mg/kg fructose or an equivalent volume of sterile saline), followed by layered suturing of the subcutaneous tissue and skin. Erythromycin ophthalmic ointment was applied topically. Sixty minutes after occlusion, the suture was gently withdrawn using ophthalmic forceps. The incision was re-sutured layer by layer, and erythromycin ophthalmic ointment was applied again.

Cerebral blood flow measurement

Under anesthesia, the mouse skull was exposed, and hair and body fluids were wiped off with normal saline. Cerebral blood flow was monitored using a laser speckle imager. After measurement, the skin was sutured, followed by topical application of cocaine paste and erythromycin ophthalmic ointment.

Mouse MRS and T2 MRI examinations

For MRS sequences, Mice were anesthetized with isoflurane. Their body temperature was maintained at 37 °C, and their respiratory rate was controlled at 100-120 breaths per minute. Scanning was performed on a uMR 9.4 T system with a voxel size of 2.0 × 1.5 × 2.0 mm. Strong water suppression was applied with a suppression bandwidth of 96 Hz. For T2 sequences, Mice were anesthetized with isoflurane, and their body temperature was maintained at 37 °C with a respiratory rate of 100-120 breaths per minute. Scanning was conducted using a uMR 9.4 T system. The voxel size was 0.10 × 0.10 × 0.50 mm, with a slice thickness of 0.5 mm. A total of 30 slices were acquired for the transverse, sagittal, and coronal planes, respectively.

TTC Staining

Mice were anesthetized with isoflurane and perfused transcardially with prewarmed normal saline at 37 °C. Brain tissues were harvested and cut into slices of 1.5 mm thickness. The sections were immediately immersed in 2% TTC solution, incubated at 37 °C for 10 min, and then photographed.

Blood-Brain Barrier Permeability Assay

Mice were injected intravenously via the tail vein with 0.5% Evans blue (EB) at a dose of 0.1 mL per 10 g body weight. One hour later, the animals were anesthetized and transcardially perfused with normal saline to flush out intravascular dye. Brains were isolated and cut into 1.5 mm-thick slices, which were photographed immediately afterwards.

Mouse electroencephalography (EEG) monitoring

Mice were implanted with electrodes on the head and allowed to recover for one week before tMCAO surgery. Electrical signals from the contralateral and injured hemispheres were recorded before, during, and after ischemia. Six frequency bands were defined: δ, θ, α, β, γ, and high γ. The power spectral density (PSD) was calculated using Welch's method. Power values within each band were summed, and their relative proportions to the total power were computed. The time window was set to 1 minute, and recordings of 20-30 minutes were selected for each phase.

Animal Metabolic Flux and Metabolomics Sample Preparation

For metabolomic analysis, mice subjected to the tMCAO model were deeply anesthetized at different time points after the onset of ischemia. The animals were transcardially perfused with ice-cold normal saline. Brain tissues were rapidly dissected, placed into centrifuge tubes, and immediately snap-frozen in liquid nitrogen. For metabolic flux analysis, mice were intraperitoneally injected with isotopically labeled glucose or fructose at a dose of 93 mg/kg immediately after filament insertion during tMCAO surgery. Two hours later, the mice were deeply anesthetized and transcardially perfused with ice-cold normal saline. Brain tissues were rapidly dissected, transferred into centrifuge tubes, and snap-frozen in liquid nitrogen. Subsequently, the extraction solution (acetonitrile:methanol:water = 4:4:2) was added, and the samples were homogenized on ice. The homogenates were stored at −80 °C for at least 2 h, then centrifuged at 12,000 rcf for 15 min at 4 °C. The supernatants were collected into pre-chilled centrifuge tubes for subsequent LC/MS analysis. For NADH quantification, calibration standards were prepared in the same extraction solution and analyzed via LC/MS.

Brain mass spectrometry imaging (MSI)

Mice subjected to tMCAO surgery were deeply anesthetized and transcardially perfused with ice‑cold normal saline. Brain tissues were rapidly dissected, wrapped in aluminum foil, and frozen on liquid nitrogen to avoid direct liquid nitrogen contact. To precisely characterize regional metabolic alterations within ischemic lesions, metabolite profiles in brain tissues were analyzed using matrix‑assisted laser desorption/ionization (MALDI) mass spectrometry imaging (MSI). Frozen brain tissues were embedded in 10% gelatin for cryosectioning and subsequently subjected to MSI analysis to profile glycolytic intermediates, TCA cycle metabolites, and NADH. For absolute NADH quantification, serial standard solutions were prepared in 10% gelatin, sectioned under identical conditions, and analyzed alongside brain tissue sections via MSI. For the detection of [U‑¹³C₆] ‑fructose‑labeled metabolites, 3 mg/mL 1,5‑diaminonaphthalene (1,5‑DAN) matrix solution was prepared using methanol: acetonitrile: water (3:3:2, v/v/v) and uniformly sprayed onto tissue sections. Matrix‑coated sections were mounted on target plates for MSI acquisition. Tissue regions were visualized and focused using the AP SMALDI imaging system software. Briefly, the Z‑axis was adjusted to obtain clear tissue visualization, the X and Y axes were used to define the scanning region, and fine Z‑axis tuning was performed to optimize laser spot focusing and signal intensity. Mass spectral data were acquired via pixel‑by‑pixel laser scanning. Laser irradiation induced desorption and ionization of analytes co‑crystallized with the matrix, and generated ions were delivered to the mass spectrometer to obtain spatial m/z information across the entire tissue section.

Two-photon imaging in mice

Mice were intravenously injected with SoNar, mito-ROS, and GO-ATeam2 probes to detect the NAD⁺/NADH ratio, mitochondrial ROS, and ATP levels in neurons within the cerebral lesion area, respectively. Two weeks after injection, cranial window surgery was performed, and a 5 mm-diameter glass coverslip was implanted. The mice were allowed to recover for another two weeks. Baseline signals were acquired before tMCAO surgery. During ischemia, signals at the identical position in the lesion region were continuously recorded. Filaments were withdrawn to restore blood flow 60 minutes later, and signals were further monitored at the same site after reperfusion. Finally, the signals obtained during ischemia and reperfusion were normalized to the baseline values.

Electron Microscopy Sample Preparation

For cell samples, the original culture medium was aspirated, and cells were rinsed twice with HBSS. Then, a pre-warmed 2.5% glutaraldehyde solution at 37 °C was added for fixation. Fixed samples were stored at 4 °C for subsequent electron microscopy sample preparation.

For tissue samples, mice were deeply anesthetized and transcardially perfused with PB buffer. Next, 10 mL of pre-warmed 2.5% glutaraldehyde solution (37 °C) was delivered via perfusion, followed by 40 mL of ice-cold 2.5% glutaraldehyde solution (4 °C). Brain tissues were rapidly harvested and immersed in 2.5% glutaraldehyde solution. Target regions were sectioned using a vibratome, and sections were kept at 4 °C before electron microscopy processing.

Quantitative analysis using Fiji

Fiji software was used to quantify fluorescence intensity and neuronal area from confocal images following standard protocols to ensure reliable results. For fluorescence intensity analysis, regions of interest (ROIs) were selected on target neurons. Target areas were identified via thresholding, with identical threshold values applied across all groups. The Analyze function was used to record the mean and integrated fluorescence intensity of each ROI. At least 10 individual neurons were measured per sample, and the mean was used for statistical analysis. For neuronal area quantification, images from neuron-specific fluorescence channels were used. ROIs and threshold settings were consistent among groups, and measurements were performed only on threshold-defined regions. A minimum of 5 independent fields of view were examined for each sample.

Data Statistical Analysis Methods

All statistical analyses and figure plotting were performed with GraphPad Prism 9.0 to ensure standardized, accurate, and reproducible data presentation. Quantitative results are presented as mean ± standard deviation (SD) or mean ± standard error of the mean (SEM), as indicated in figure legends.

The Shapiro-Wilk test was applied to verify normal data distribution, and Levene’s test was used to evaluate homogeneity of variance among groups.

For two-group comparisons, unpaired Student’s t-test was adopted for normally distributed data with equal variance; the nonparametric Mann-Whitney U test was utilized for datasets that violated normality or homogeneity of variance assumptions.

Multiple comparisons involving a single grouping factor were analyzed via one-way analysis of variance (ANOVA): Dunnett’s post-hoc test was used when all experimental groups were compared against a unified control group, and Tukey’s HSD post-hoc test was applied for all pairwise comparisons between groups. Two-way analysis of variance (ANOVA) was conducted to evaluate two independent variables and their interaction, followed by Sidak’s multiple comparisons test for simple effect pairwise analyses.

Non-normally distributed data were analyzed using the Kruskal-Wallis H test, with Dunn’s multiple comparison test as the corresponding post-hoc test.

The threshold for statistical significance was set at α = 0.05; a P value < 0.05 was regarded as statistically significant. All bar and line graphs were generated within GraphPad Prism 9.0.

Clinical trial

This single-center, prospective, randomized, open-label, blinded-endpoint (PROBE) exploratory phase IIa trial was registered on ClinicalTrials.gov (NCT07626541). We planned to enroll 46 participants equally allocated to two arms. Eligible patients presented with acute ischemic stroke within 4.5 hours of symptom onset and qualified for intravenous thrombolysis. Forty-four patients were included in the intention-to-treat (ITT) analysis, with 21 assigned to the fructose intervention group and 23 to the control group. The intervention group received a single 250 mL 10% fructose solution initiated prior to thrombolysis, with the 60-minute infusion administered without delaying reperfusion therapy; all participants received standardized intravenous thrombolysis and routine stroke ward care. Serial neurological assessments were performed using the National Institutes of Health Stroke Scale (NIHSS) at prespecified time points. The relative NIHSS change at 72 hours post-thrombolysis was defined as the prespecified primary endpoint. Secondary exploratory endpoints included 7-day and 30-day NIHSS scores, 24-hour post-recanalization cerebral MRI metrics, ¹H-MRS cerebral metabolite quantification, and 30-day modified Rankin Scale (mRS) functional outcomes.

Structural MRI including DWI and T2-FLAIR was acquired solely at 24 hours after thrombolysis. No pre-thrombolysis baseline DWI scans were obtained, so only single-time-point 24 h infarct volume and cerebral edema were quantified. ¹H-MRS scans were collected at the identical 24-hour window to measure cerebral NAA, choline and lactate concentrations. Long-term functional recovery was evaluated via mRS at the 30-day follow-up. Safety surveillance throughout the 30-day follow-up covered intracranial hemorrhage, hepatorenal dysfunction, glycemic fluctuation, and neurological deterioration. Venous blood glucose was tested immediately upon fructose infusion completion and at the subsequent fasting morning. All adverse events were graded per CTCAE v5.0, with serious adverse events documented in real time. Predefined subgroup analyses stratified by baseline clinical characteristics were performed. All imaging and scale assessments were adjudicated by a treatment-blinded clinical committee to reduce evaluation bias.

All statistical analyses were conducted in GraphPad Prism. Non-normal continuous variables were compared via the Mann-Whitney U test; categorical variables were analyzed using Chi-squared or Fisher’s exact test as appropriate. Repeated-measures ANOVA was used for longitudinal NIHSS measurements. All tests were two-sided with α = 0.05. No multiple comparison correction was implemented for secondary endpoints given the exploratory phase IIa design and limited sample size; significant secondary findings are interpreted exclusively as hypothesis-generating signals rather than definitive therapeutic evidence.

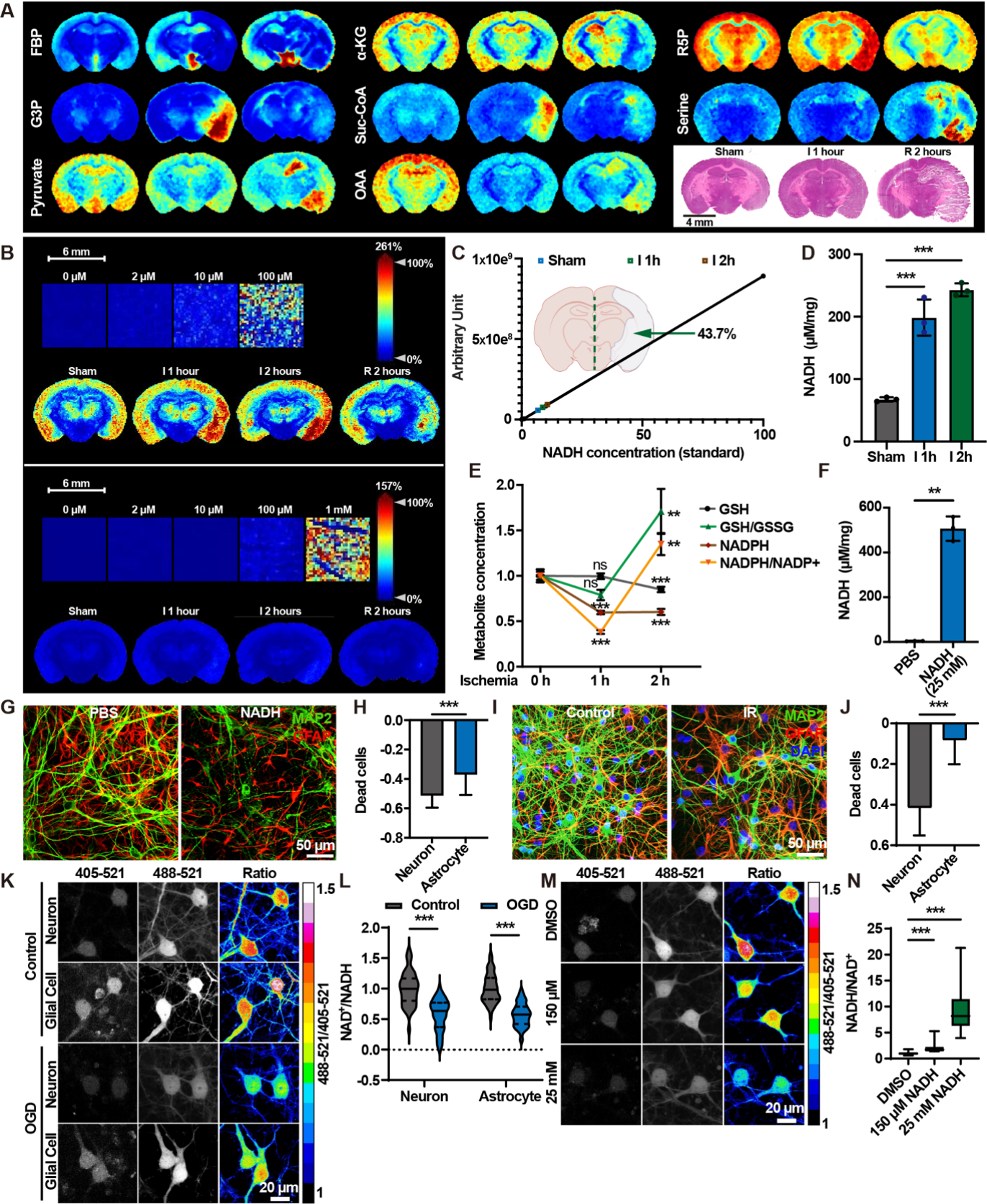

Fig. S1. Neurons display elevated vulnerability to disruptions in NAD(H) redox homeostasis

**(A)** Mass spectrometry imaging reveals metabolite alterations in the injured brain regions of mice at distinct stages of ischemia-reperfusion.

**(B-D)** Mass spectrometry imaging was applied to map the spatial distribution of cerebral NADH pools, and parallel LC-MS analysis enabled precise absolute quantification of NADH concentrations in damaged brain tissue (N=3).

**(E)** LC-MS metabolomic profiling measured tissue concentrations of GSH, the GSH/GSSG ratio, NADPH, and the NADPH/NADP⁺ ratio in lesioned mouse brain tissue harvested across sequential ischemic time points (N=3).

**(F)** LC-MS-based absolute quantification of intracellular NADH levels in mixed neuron-astrocyte co-cultures following exogenous NADH supplementation (N=3).

**(G and H)** Mixed primary neuron-astrocyte co-cultures were incubated with 25 mM NADH for 4 h to induce reductive stress, followed by viability quantification to compare cell-type-specific survival outcomes (N=4, n>5).

**(I and J)** Mixed primary neuron-astrocyte co-cultures were subjected to 4 h OGD. Cell survival was assessed via dual immunolabeling for the neuronal marker MAP2 and astrocytic marker GFAP, revealing divergent viability responses between the two cell types under hypoxic insult (N=4, n>5).

**(K and L)** Primary neurons and glia were transduced with the SoNar redox biosensor to measure basal NAD⁺/NADH ratios. After 4 h OGD, cell-type-specific shifts in the NAD⁺/NADH redox ratio were quantified via ratiometric fluorescence imaging (N=4, n>5).

**(M and N)** Primary neurons expressing the SoNar probe were treated with PBS vehicle, 150 µM NADH, or 25 mM NADH, and subsequent ratiometric imaging captured dose-dependent changes in intracellular NAD⁺/NADH ratios (N=4, n>5).

Data are presented as means ± SD. Multiple-group comparisons were performed using one-way analysis of variance (ANOVA), followed by Dunnett’s post-hoc test (D, E and N); two-way analysis of variance (ANOVA) was performed, followed by Sidak’s multiple comparisons test (L); comparisons between two groups were conducted using the unpaired Student’s t-test (F, H and J). *P < 0.05, **P < 0.01, ***P < 0.001; ns, not significant.

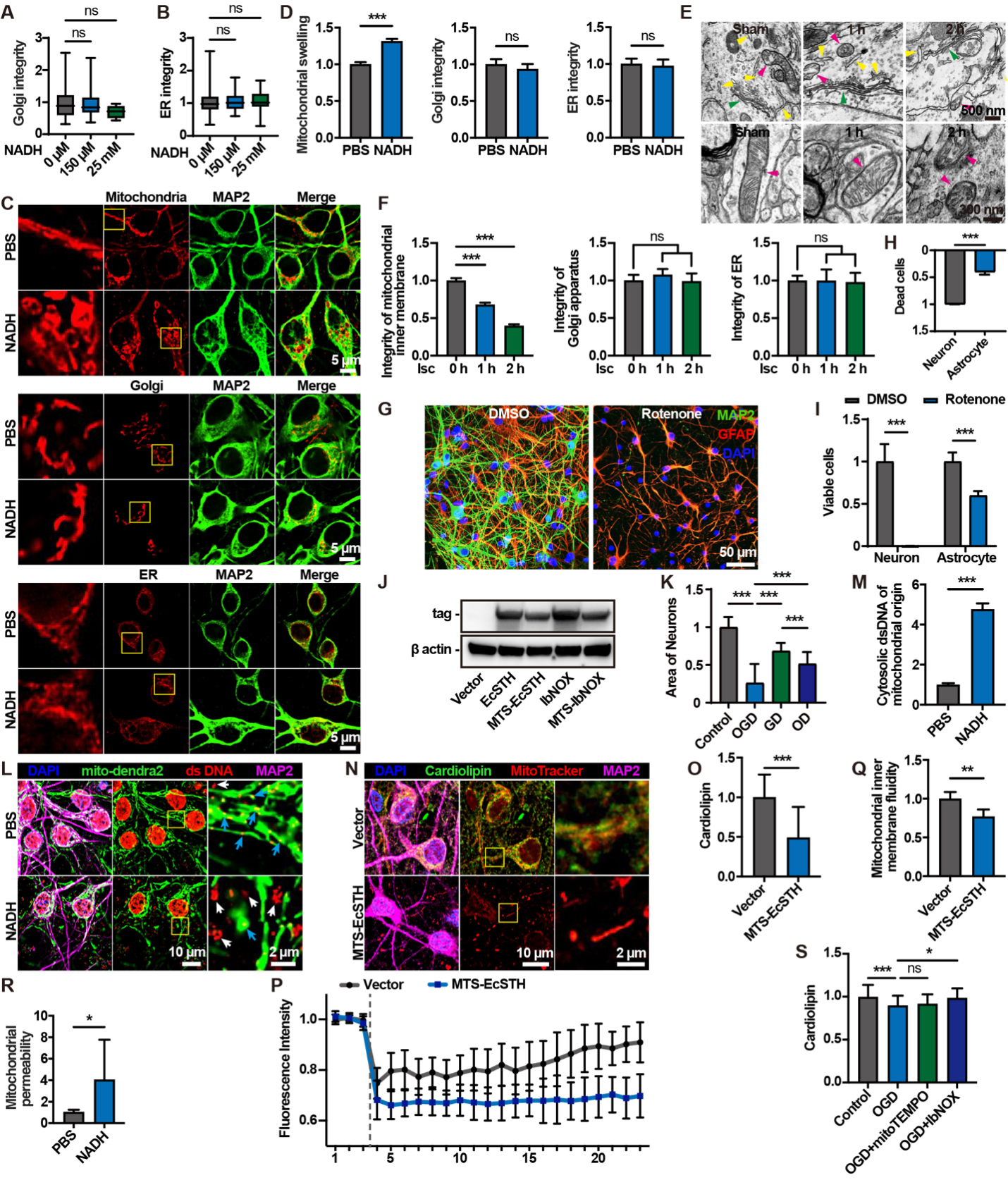

Fig. S2. Reductive stress preferentially damages neuronal mitochondria

**(A and B)** Primary neurons were incubated with either 150 µM or 25 mM NADH, followed by transmission electron microscopy (TEM) imaging and quantitative scoring of Golgi apparatus (A) (N=5, n>1) and endoplasmic reticulum (ER) structural integrity (B) (N=4, n>5).

**(C and D)** Primary neurons were transduced with fluorescent reporters for mitochondria and ER, or immunolabeled for the Golgi complex. Following 4 h incubation with 25 mM NADH, confocal microscopy was performed to assess structural integrity of mitochondria, Golgi, and ER compartments (N=5, n>6).

**(E and F)** Mouse brain tissue from sham-operated animals and mice subjected to 1 h or 2 h ischemia was processed for TEM imaging. Quantitative analysis was conducted to evaluate the structural integrity of mitochondria, Golgi apparatus, and ER across experimental groups (N=4, n>2).

**(G-I)** Mixed primary neuron-astrocyte co-cultures were treated with 1 µM rotenone (Rot) for 4 h to induce respiratory chain suppression, after which cellular viability was quantified to compare cell-type-specific survival (N=4, n>5).

**(J)** Western blot (WB) analysis to characterize protein expression of EcSTH, MTS-EcSTH, lbNOX, and MTS-lbNOX in primary neuronal cultures.

**(K)** Primary neurons were subjected to 24 h OGD, glucose deprivation alone, or oxygen deprivation alone; MAP2 immunostaining was used to quantify neuronal viability under each stress (N=6, n>6).

**(L and M)** Primary neurons pre-labeled with the mitochondrial reporter mito-dendra2 were treated with PBS vehicle or 10 mM NADH for 4 h. Immunofluorescence was used to quantify mitochondrial DNA release into the cytosol (N=4, n>5).

**(N and O)** Primary neurons were transduced with MTS-EcSTH to elevate intramitochondrial NADH pools. Mitochondria were co-stained with the cardiolipin-specific dye NAO and the mitochondrial membrane tracker CMTMRos, and cardiolipin abundance was quantified (N=4, n>5).

**(P and Q)** Mitochondrial membranes of primary neurons were labeled with CMTMRos. Fluorescence recovery after photobleaching (FRAP) of small mitochondrial subdomains was measured to compare mitochondrial membrane fluidity between neurons transfected with empty vector or MTS-EcSTH (N>4).

**(R)** Mitochondrial permeability was assayed in primary neurons following 4 h incubation with 25 mM NADH (N=3, n>3).

**(S)** Primary neurons transfected with mito-lbNOX (empty vector as control) or treated with mitoTEMPO (DMSO as control) were subjected to 4 h of OGD. Post-OGD co-staining with NAO and CMTMRos enabled quantitative measurement of cardiolipin content (N=4, n>5).

Data are presented as means ± SD. Multiple-group comparisons were performed using one-way analysis of variance (ANOVA), followed by Dunnett’s post-hoc test (A, B, F and S) or Tukey’s HSD post-hoc test (I and K); Comparisons between two groups were conducted using the unpaired Student’s t-test (D, H, M, O, Q and R). *P < 0.05, **P < 0.01, ***P < 0.001; ns, not significant.

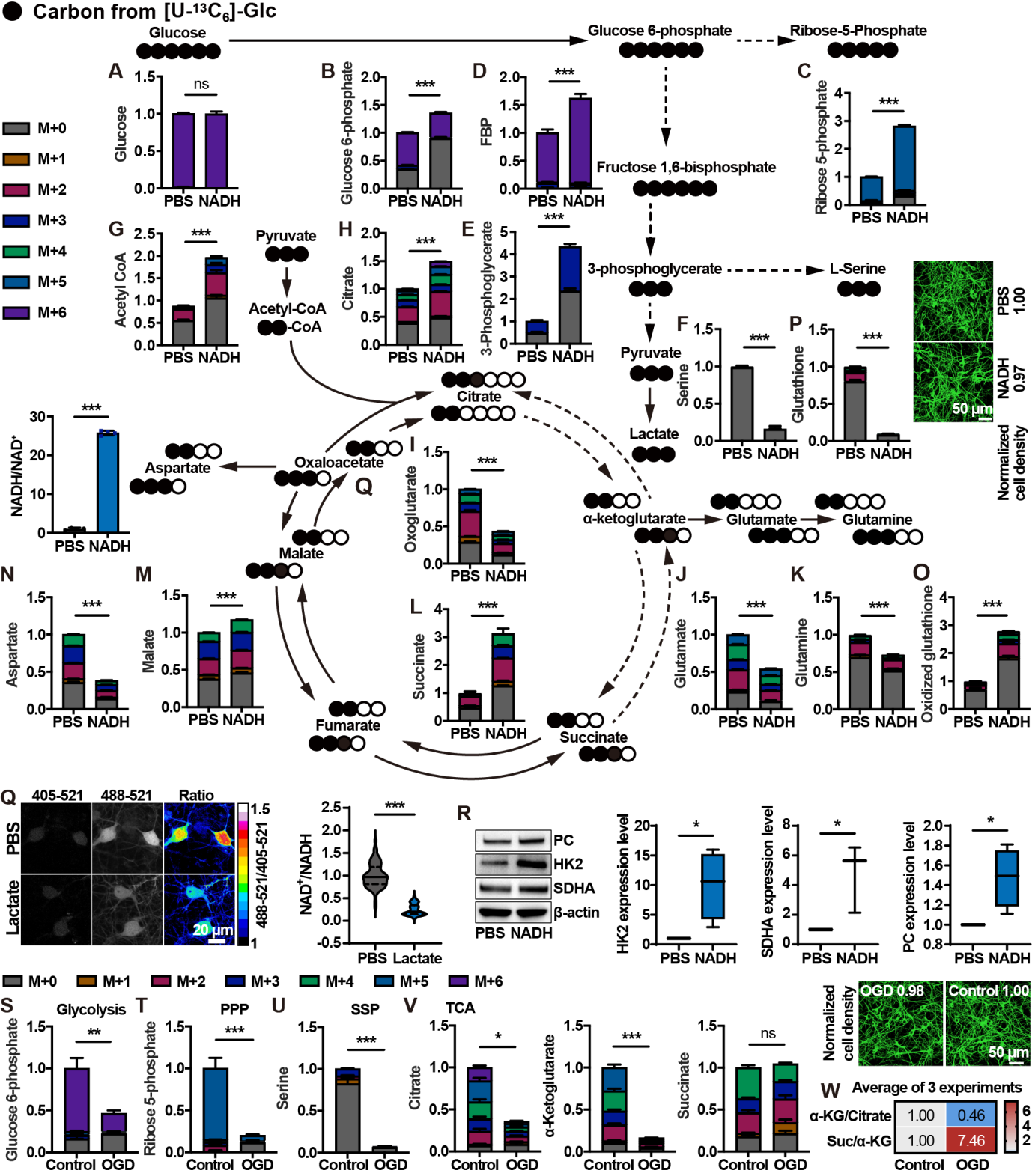

Fig. S3. Reductive stress reshapes core metabolic flux in neurons

**(A-P)** ¹³C-isotope tracing metabolomics was performed to systematically profile metabolic remodeling triggered by exogenous NADH in primary neurons (N=4).

**(Q)** Primary neurons expressing the SoNar redox biosensor were incubated with PBS vehicle or 10 mM sodium lactate. Ratiometric fluorescence quantification of SoNar signals was conducted to monitor dynamic shifts in mitochondrial redox status (N=4, n>5).

**(R)** Western blot (WB) quantification of HK2 (hexokinase 2) (N=4), SDHA (succinate dehydrogenase complex subunit A) (N=3), and PC (pyruvate carboxylase) (N=4) protein abundance in NADH-treated neurons to assess transcriptional and post-translational regulation of central metabolic cascades.

**(S-U)** ¹³C-glucose tracing assays measured abundance shifts of glucose-6-phosphate (G6P) (a rate-limiting glycolytic intermediate) (S), ribose-5-phosphate (R5P) (the canonical pentose phosphate pathway marker metabolite) (T), and serine (the principal output of the serine biosynthetic pathway) (U), under OGD conditions (N=3).

**(V)** ¹³C-glucose tracing was implemented to profile alterations in canonical TCA cycle intermediates including citrate, α-ketoglutarate (α-KG), and succinate upon OGD treatment (N=3).

**(W)** Molar ratios of isotopically labeled TCA-cycle metabolites relative to their upstream precursors were calculated to infer relative pathway flux rates (N=3).

Data are presented as means ± SD. Comparisons between two groups were conducted using the unpaired Student’s t-test. *P < 0.05, **P < 0.01, ***P < 0.001; ns, not significant.

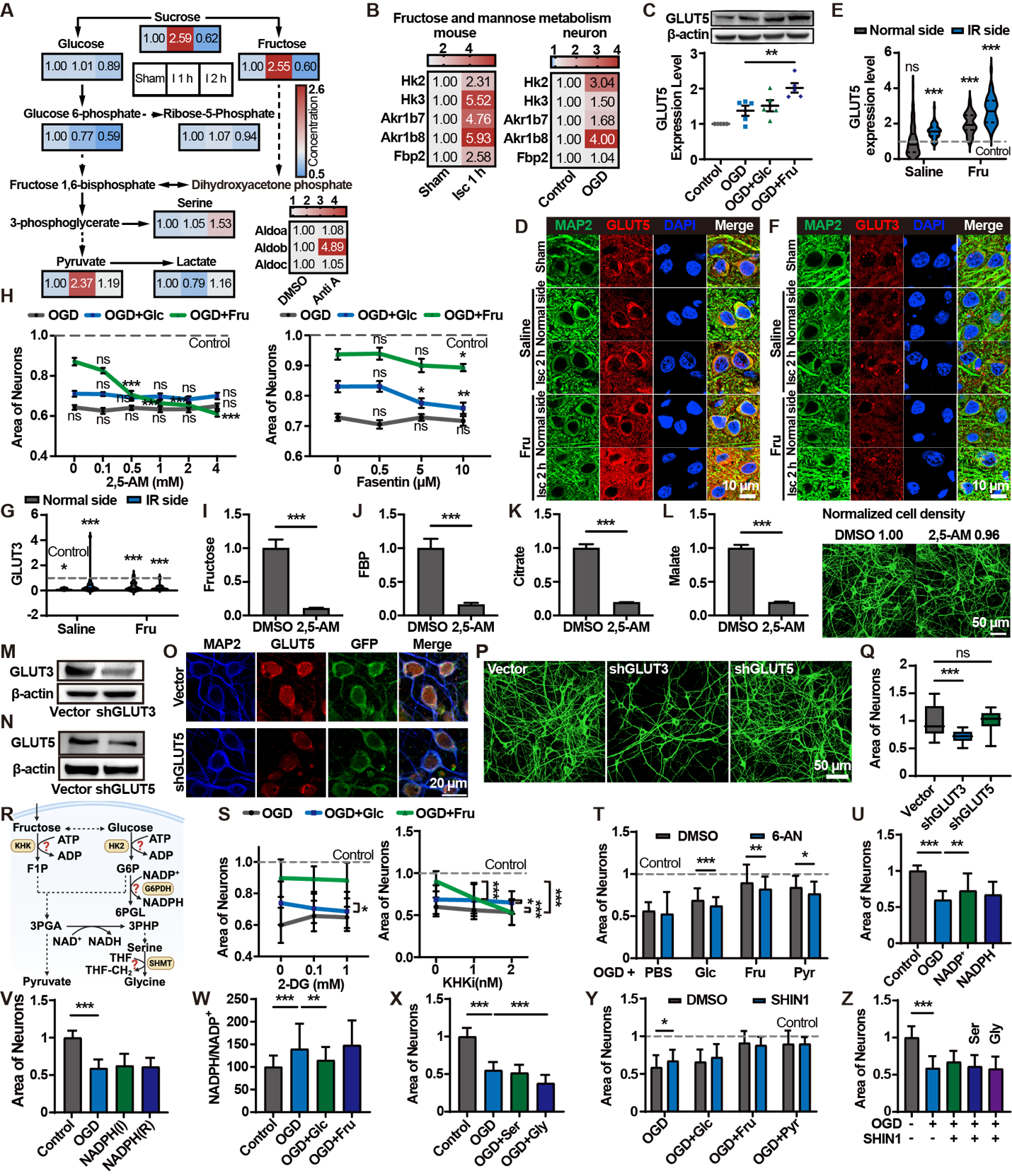

Fig. S4. Neurons express GLUT5, and GLUT5 inhibition abrogates fructose-mediated neuroprotection

**(A)** Mouse brain tissue and primary neurons were harvested and processed for metabolomic profiling (mouse brain tissue, n=3) and RNA-seq (primary neurons, n=4) to assess changes in metabolites and transcripts linked to metabolic pathways.

**(B)** RNA-seq was performed on brain tissue from sham-operated mice and mice subjected to 1 h ischemia (N=5), as well as primary neurons under normoxic control or OGD conditions (N=4). Differential expression of core genes governing mannose and fructose metabolism was quantified across all groups.

**(C)** Western blot (WB) analysis of GLUT5 protein abundance in primary neurons under distinct experimental conditions (N=5).

**(D and E)** Mice were subjected to 2 h permanent cerebral ischemia and left untreated or administered 90 mg/kg fructose. Dual immunohistochemical staining for GLUT5 and the neuronal marker MAP2 was conducted to quantify neuronal GLUT5 expression (N=4, n>5).

**(F and G)** Following 2 h permanent cerebral ischemia, mice received vehicle or 90 mg/kg fructose. Co-immunolabeling of GLUT3 and MAP2 was used to quantify GLUT3 protein levels within cortical neurons (N=4, n>5).

**(H)** Primary neurons were subjected to OGD-reperfusion injury, with glucose or fructose supplemented during OGD. Neurons under OGD were co-treated with either the selective GLUT5 inhibitor 2,5-anhydro-D-mannitol (2,5-AM) (N=4, n>5) or the GLUT1-specific inhibitor fasentin (N=4, n>5).

**(I-L)** Primary neurons were exposed to 4 h OGD alongside 10 mM isotopically labeled fructose, with parallel delivery of DMSO or 2,5-anhydro-D-mannitol (2,5-AM). LC-MS metabolic tracing quantified isotopic flux distribution throughout central carbon metabolism. Data are presented as mean ± SD from three independent biological replicates (n = 4 culture replicates per group). Corresponding MAP2 immunofluorescence images confirmed equivalent neuronal cell densities between groups at the time of LC-MS sample collection (N=4).

**(M)** Primary neurons were transfected with empty control vector or GLUT3-targeting shRNA (shGLUT3), and GLUT3 knockdown efficiency was validated via Western blot.

**(N)** Primary neurons were transduced with empty vector or shRNA against GLUT5 (shGLUT5); Western blot analysis verified the degree of GLUT5 protein suppression.

**(O)** Primary neurons transfected with shGLUT5 or empty vector were co-stained for GLUT5 (red) and MAP2 (green). GFP signal carried by the lentiviral vector (green) demarcated successfully transfected neurons, enabling quantification of GLUT5 abundance within target cells.

**(P and Q)** Primary neurons were transfected with empty vector, shGLUT5, or shGLUT3 under basal culture conditions. MAP2 immunofluorescence was used to quantify neuronal viability for each genetic manipulation group (N=4, n>5).

**(R)** Schematic illustrating two routes for fructose entry into glycolysis via ketohexokinase (KHK) or hexokinase (HK), with downstream branching toward the pentose phosphate pathway (PPP) or serine biosynthetic pathway.

**(S)** Primary neurons subjected to OGD-reperfusion received glucose or fructose during OGD, alongside DMSO, 2-DG, or the KHK inhibitor (KHKi). Neuronal survival was quantified via MAP2 staining (N=4, n>5).

**(T)** During OGD-reperfusion, neurons were supplemented with glucose or fructose plus vehicle DMSO or 100 nM 6-aminonicotinamide (6-AN). MAP2 immunostaining was used to assess neuronal viability (N=4, n>5).

**(U)** Primary neurons underwent OGD-reperfusion with PBS, 1 µM NADP⁺, or 10 µM NADPH provided throughout OGD; MAP2 staining quantified neuronal survival rates (N=4, n>5).

**(V)** 10 µM NADPH was supplemented exclusively during the ischemic phase or sustained through both ischemia and reperfusion in OGD-reperfused neuronal cultures. MAP2 immunostaining was performed to compare neuronal survival across treatment schedules (N=4, n>5).

**(W)** Primary neurons expressing the NADPH/NADP⁺ biosensor iNAP were subjected to 4 h OGD, and ratiometric imaging quantified shifts in the NADPH/NADP⁺ redox ratio (N=4, n>5).

**(X)** Neuronal cultures received PBS, 0.4 mM L-serine, or 0.8 mM glycine during OGD-reperfusion, followed by MAP2 staining to quantify survival (N=4, n>5).

**(Y)** Primary neurons were exposed to OGD-reperfusion with glucose or fructose co-administered alongside DMSO or 10 µM SHIN1, a selective PHGDH inhibitor. MAP2 immunofluorescence was used to evaluate neuronal viability (N=4, n>5).

**(Z)** OGD-reperfused neurons received serine or glycine during ischemia, with concurrent treatment of DMSO or 10 µM PHGDH inhibitor SHIN1. MAP2 staining was performed to quantify differences in neuronal survival (N=4, n>5).

Data are presented as means ± SD. Multiple-group comparisons were performed using one-way analysis of variance (ANOVA), followed by Dunnett’s post-hoc test (C, H, Q, S, U, V, W, X and Z); two-way analysis of variance (ANOVA) was performed, followed by Sidak’s multiple comparisons test (E, G, T and Y); comparisons between two groups were conducted using the unpaired Student’s t-test (I, J, K and L). *P < 0.05, **P < 0.01, ***P < 0.001; ns, not significant.

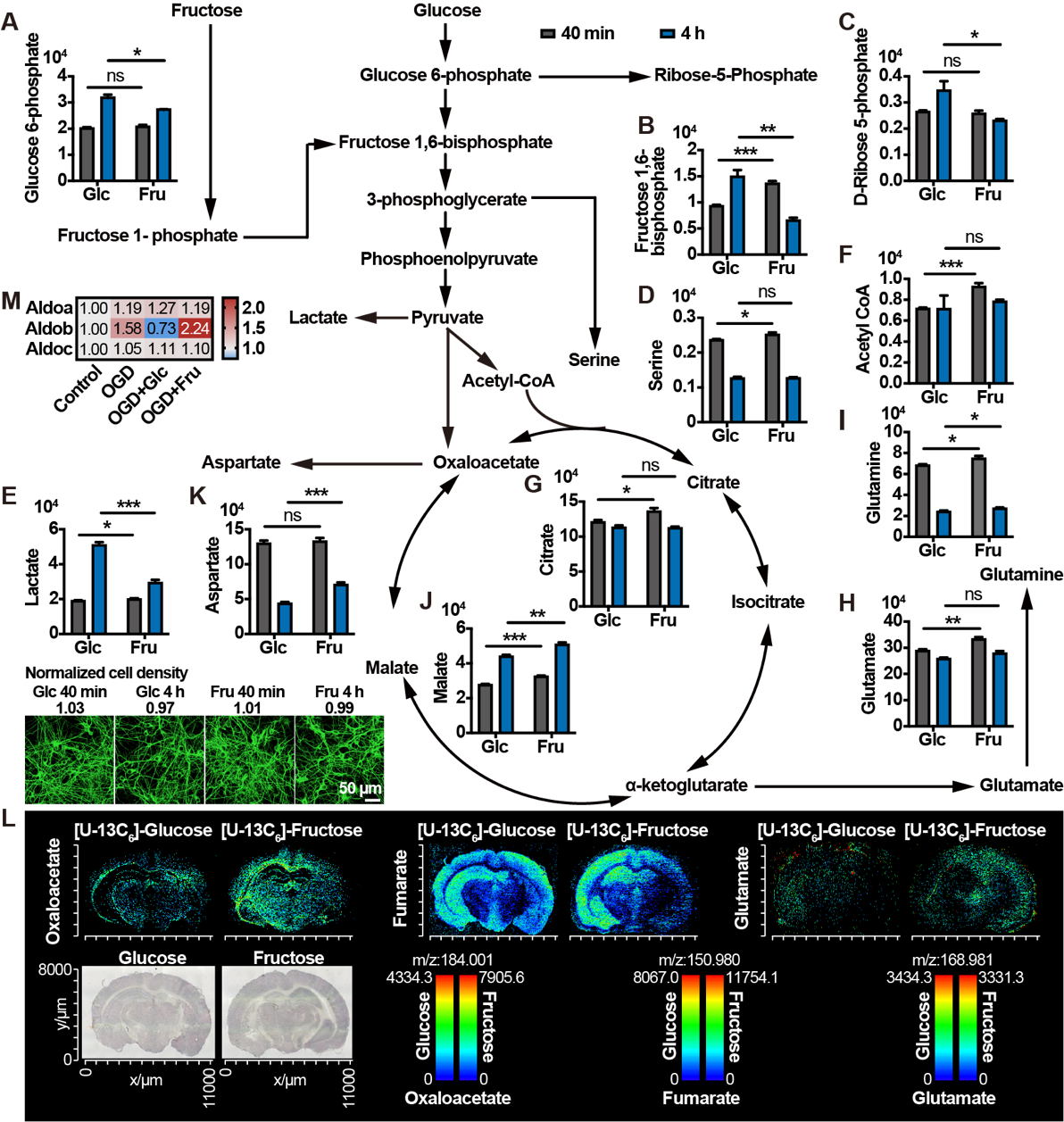

Fig. S5. Neurons catabolize fructose far more robustly than glucose

**(A-K)** Primary neurons were cultured with 10 mM glucose or 10 mM fructose during 40 min or 4 h OGD. LC-MS metabolomic profiling uncovered divergent intracellular metabolite signatures induced by the two carbon substrates under hypoxic deprivation. Corresponding MAP2 immunofluorescence images confirmed equivalent neuronal cell densities between groups prior to LC-MS metabolite extraction (N=4).

**(L)** At the onset of tMCAO occlusion, mice received an intraperitoneal bolus of 93 mg/kg [U-¹³C₆] -glucose and [U-¹³C₆]-fructose. Brain tissue was harvested 1 h post-ischemia for mass spectrometry imaging to map the spatial distribution of isotope-incorporated metabolites.

**(M)** RNA-seq was performed on primary neurons subjected to distinct culture treatments to quantify differential mRNA expression of Aldoa, Aldob and Aldoc across experimental cohorts (N=4).

Data are presented as means ± SEM. Multiple-group comparisons were performed using one-way analysis of variance (ANOVA), followed by Tukey’s HSD post-hoc test. *P < 0.05, **P < 0.01, ***P < 0.001; ns, not significant.

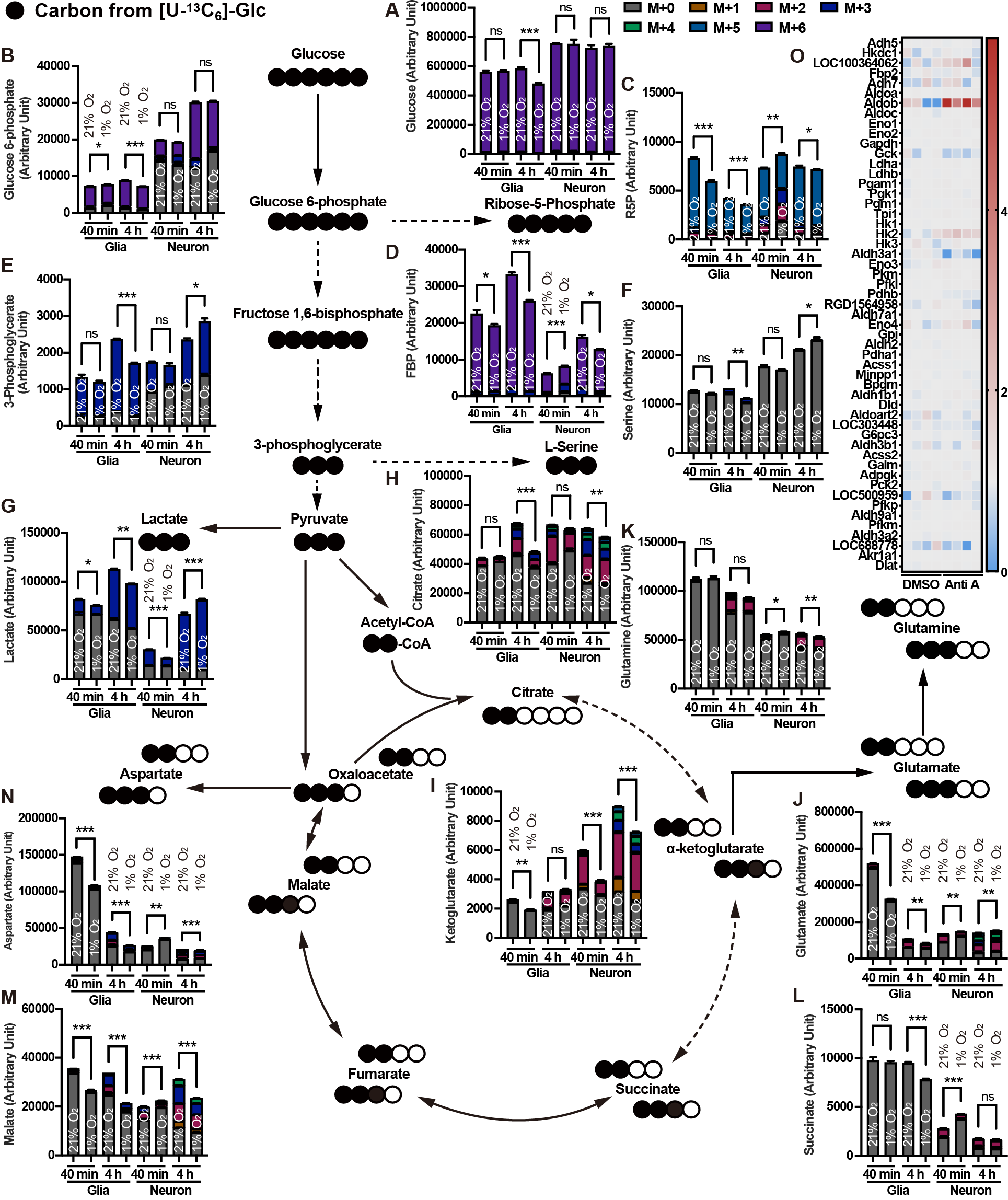

Fig. S6. Neurons retain functional mitochondrial metabolic turnover under hypoxic conditions

**(A-N)** Primary astrocytes and primary neurons were exposed to 40 min or 4 h OGD, and ¹³C-isotope tracing metabolomics was applied to profile cell-type-specific metabolic remodeling under hypoxic deprivation (N=3).

**(O)** Primary neurons were incubated with vehicle DMSO or antimycin A for 4 h, followed by RNA-seq transcriptional profiling to quantify shifts in the expression of core glycolytic genes (N=4).

Data are presented as means ± SD. Two-way analysis of variance (ANOVA) was performed, followed by Sidak’s multiple comparisons test. *P < 0.05, **P < 0.01, ***P < 0.001; ns, not significant.

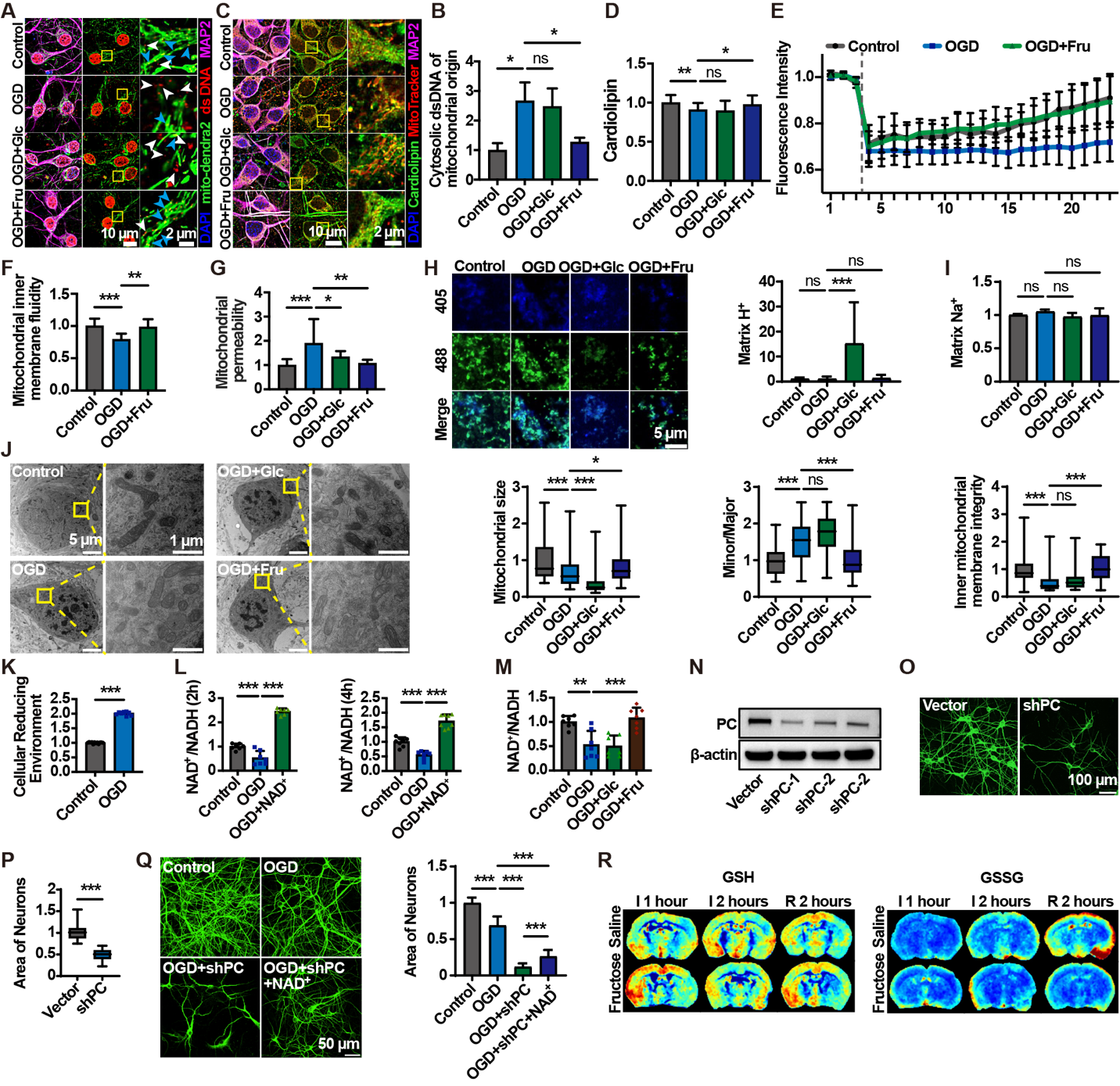

Fig. S7. Fructose treatment preserves mitochondrial homeostasis

**(A and B)** Primary neurons were subjected to 4 h OGD with glucose or fructose supplementation, and mitochondrial DNA release into the cytosol was quantified (N=4, n>5).

**(C and D)** Primary neurons underwent 4 h OGD in the presence of glucose or fructose, followed by quantitative measurement of mitochondrial cardiolipin abundance (N=4, n>5).

**(E and F)** Primary neurons were treated with fructose or PBS during OGD, and mitochondrial membrane fluidity was assessed (N=3, n>2).

**(G)** Primary neurons were exposed to OGD with glucose or fructose provided, and mitochondrial permeability was assayed (N=4, n>2).

**(H)** Isolated mitochondria harvested from primary neurons after 4 h OGD were loaded with BCECF-AM. Ratiometric fluorescence imaging with BCECF-AM was used to track dynamic proton concentrations within the mitochondrial matrix (N=5, n>5).

**(I)** Isolated mitochondria from primary neurons subjected to 4 h OGD were stained with SBFI-AM. Ratiometric measurement was performed to quantify matrix proton dynamics (N=4, n>5).

**(J)** Transmission electron microscopy (TEM) was used to evaluate mitochondrial ultrastructural integrity in primary neurons cultured with isocaloric glucose or fructose during 4 h OGD (N=5, n>=8).

**(K)** Commercial assay kits were used to measure the global reductive status in primary neurons after 4 hours of OGD treatment (N=6, n>2).

**(L)** Enzymatic cycling assays were applied to quantify intracellular NAD⁺/NADH ratios in primary neurons treated with PBS vehicle or exogenous NAD⁺ over 2 h or 4 h OGD (N>=7).

**(M)** Primary neurons received glucose or fructose as a carbon substrate during 4 h OGD, and enzymatic cycling assays were conducted to quantify steady-state NAD⁺/NADH levels (N=7).

**(N)** Lentiviral shRNA constructs targeting pyruvate carboxylase (PC) were delivered to primary neurons, and Western blot analysis validated distinct PC knockdown efficiencies across three independent shRNA sequences.

**(O and P)** Primary neurons transduced with PC-targeting shRNA were stained for MAP2, and neuronal survival was quantified (N=4, n>5).

**(Q)** shPC-transduced neurons were subjected to OGD-reperfusion injury with PBS or NAD⁺ supplemented exclusively during OGD. MAP2 immunostaining was performed to measure neuronal viability, with values normalized to normoxic shPC-expressing control neurons (N=4, n>5).

**(R)** Mass spectrometry imaging was performed to quantify spatial shifts in GSH and GSSG abundance within mouse brain tissue at sequential ischemic time points.

Data are presented as means ± SD. Multiple-group comparisons were performed using one-way analysis of variance (ANOVA), followed by Dunnett’s post-hoc test (B, D, F, G, H, I, J, L and M) or Tukey’s HSD post-hoc test (Q); comparisons between two groups were conducted using the unpaired Student’s t-test (K and P). *P < 0.05, **P < 0.01, ***P < 0.001; ns, not significant.

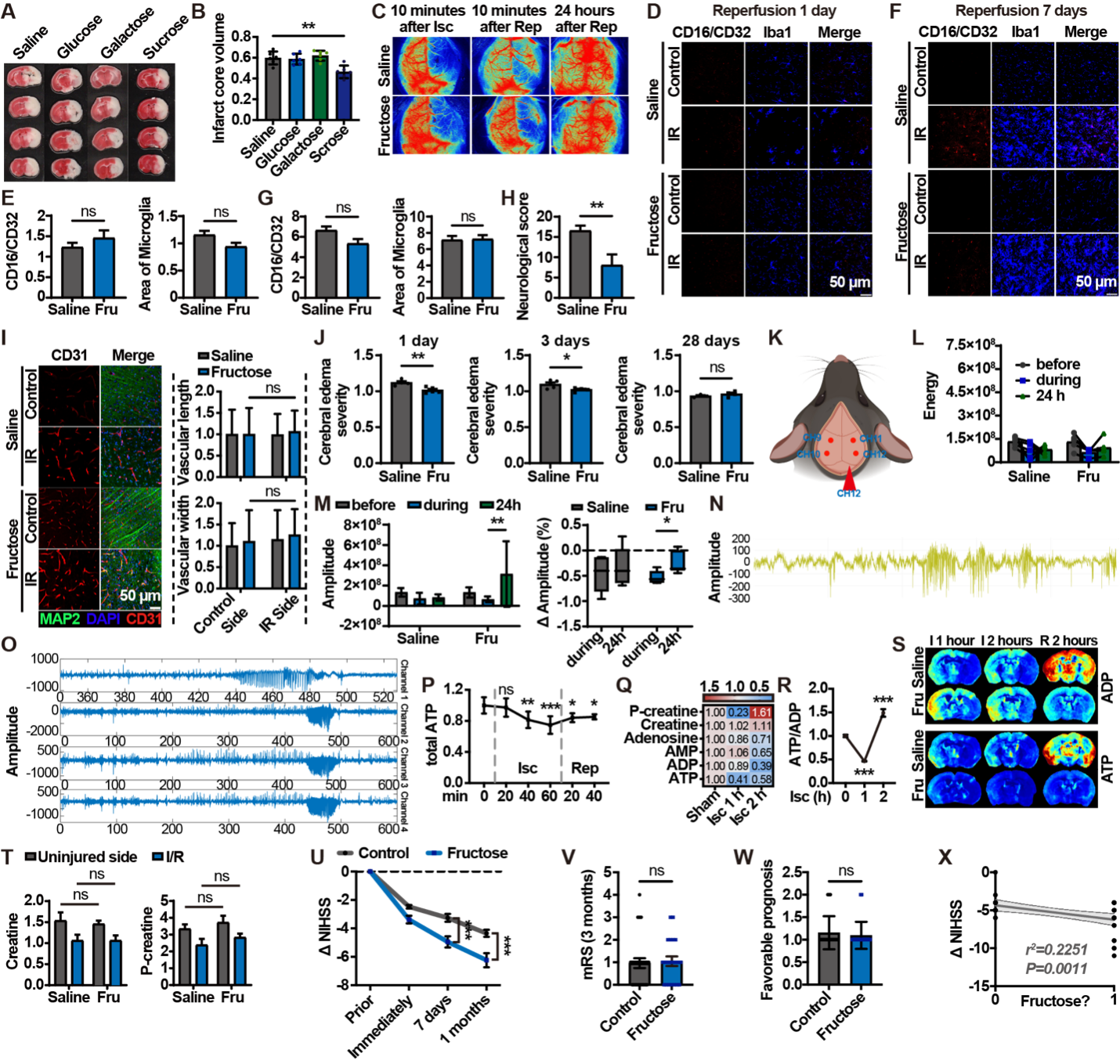

Fig. S8. Fructose exerts potent cerebral protective effects independent of cerebral blood flow and vascular remodeling, without exacerbating neuroinflammation

**(A and B)** Mice subjected to tMCAO received intravenous saline, glucose, galactose, or sucrose. TTC staining was performed to quantify cerebral infarct volume across all treatment groups (N=6; means ± SD).

**(C)** Laser speckle contrast imaging (LSCI) was used to record real-time dynamic cortical perfusion in untreated control tMCAO mice and fructose-treated tMCAO mice. Serial images were acquired at 10 min after ischemia onset, 10 min post-reperfusion, and 24 h post-reperfusion.

**(D and E)** Quantitative analysis of total microglia and pro-inflammatory activated microglia in lesioned cortical tissue at 24 h post-reperfusion (N=5, n>1; means ± SEM).

**(F and G)** Quantitative analysis of total microglia and pro-inflammatory activated microglia in lesioned cortical tissue at 7 days after reperfusion (N=5, n>1; means ± SEM).

**(H)** Neurological deficit evaluation using the Clark neurological scoring scale at 72 h post-reperfusion (N=4; means ± SD).

**(I)** Morphometric quantification of microvascular density, vessel length, and vessel width within lesioned cortical regions (N=5, n>20; means ± SD).

**(J)** Serial volumetric analysis using 9.4 T MRI at 24 h, 72 h, and 28 days post-reperfusion to compare long-term pathological progression between fructose-treated and saline control tMCAO mice (N>=4; means ± SEM).

**(K-M)** Electrodes were implanted on the skull of male C57BL/6 mice to record neuronal electrical activity. Electroencephalographic signals were captured prior to ischemia, during occlusion, and after reperfusion; signal energy, peak amplitude, and relative amplitude shift (Δ amplitude) were quantified (N=5 to 6).

**(N)** Spontaneous epileptiform discharges were detected in saline-treated mice during ischemic occlusion.

**(O)** Seizure episodes were observed in vehicle control mice under ischemic conditions.

**(P)** Primary neurons in vivo were transduced with the ATP biosensor GoAtem. Longitudinal two-photon imaging was conducted to track dynamic ATP fluctuations across sequential ischemic and reperfusion stages.

**(Q)** LC-MS metabolomic profiling quantified alterations in energy-related metabolites in brain tissue collected from sham-operated mice and animals subjected to 1 h or 2 h ischemia (N=3).

**(R)** LC-MS measurement of tissue ATP/ADP ratios in mouse brain tissue harvested at distinct ischemic time points (N=3).

**(S)** Mass spectrometry imaging with 100 μm spatial resolution was applied to tMCAO mice treated with saline or fructose. Spatial abundance gradients of ATP and ADP were quantified throughout damaged tissue, the penumbral region, and the uninjured contralateral hemisphere.

**(T)** In vivo magnetic resonance spectroscopy (MRS) scans were acquired from saline- and fructose-treated mice to quantify cerebral creatine and phosphocreatine levels within damaged brain parenchyma (N=5; means ± SEM).

**(U)** Longitudinal trends in National Institutes of Health Stroke Scale (NIHSS) scores over follow-up time points for stroke patients receiving standard treatment with or without fructose adjunct intervention.

**(V)** Modified Rankin Scale (mRS) scores assessed at 1 month after vascular recanalization in control and fructose-treated patient cohorts.

**(W)** Rates of favorable functional prognosis at 1-month post-recanalization in patients receiving saline or fructose adjuvant therapy.

**(X)** Correlation between fructose intervention and the reduction in NIHSS scores at the 1-month follow-up.

Multiple-group comparisons were performed using one-way analysis of variance (ANOVA), followed by Dunnett’s post-hoc test (B, P, and R); two-way analysis of variance (ANOVA) was performed, followed by Sidak’s multiple comparisons test (I, L, M, T and U); comparisons between two groups were conducted using the unpaired Student’s t-test (E, G, H and J). Between-group comparisons were conducted via Mann-Whitney U test (V and W). *P < 0.05, **P < 0.01, ***P < 0.001; ns, not significant.

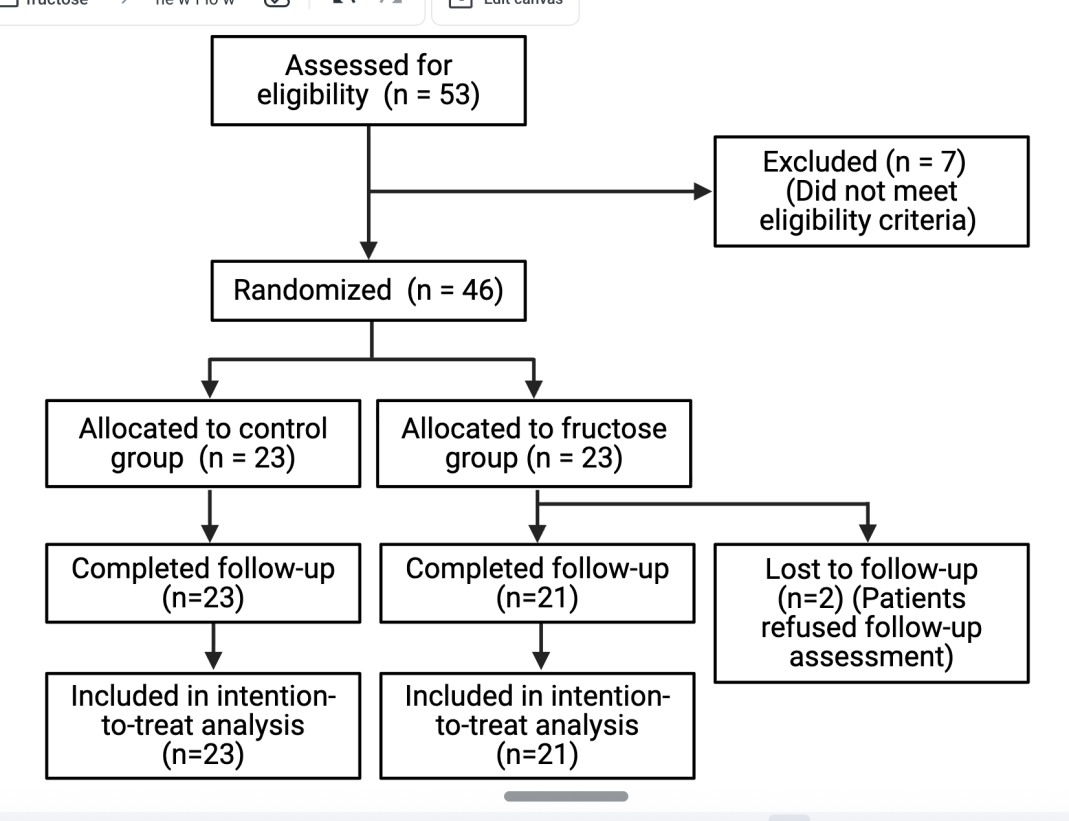

**Fig. S9. Trial profile**

Patients who had been enrolled inappropriately or who discontinued a trial drug were included in the intention-to-treat analysis, as were patients who were lost to follow-up.

| ***Indicator*** | ***Overall cohort (n=44)*** | ***Control group (n=23)*** | ***Fructose group (n=21)*** | ***P-value*** |
| --- | --- | --- | --- | --- |
| ***Demographics*** | | | | |
| *Age, median (IQR), years* | 67 (59-75) | 67 (55-74) | 68 (62-78) | 0.3001 |
| *Sex, male, n (%)* | 29 (65.9) | 15 (65.2) | 14 (66.7) | >0.9999 |
| ***Medical history*** |  | | | |
| *Diabetes mellitus, n (%)* | 5 (11.4) | 3 (13.0) | 2 (9.5) | >0.9999 |
| *Hypertension, n (%)* | 26 (59.1) | 14 (60.9) | 12 (57.1) | >0.9999 |
| *Atrial fibrillation, n (%)* | 4 (9.1) | 2 (8.7) | 2 (9.5) | >0.9999 |
| *Previous stroke/TIA, n (%)* | 11 (25.0) | 6 (26.1) | 5 (23.8) | >0.9999 |
| *Dyslipidemia, n (%)* | 22 (50.0) | 12 (52.2) | 10 (47.6) | >0.9999 |
| ***Clinical status*** | | | | |
| *Baseline NIHSS (pre-thrombolysis), median (IQR)* | 5 (5-6) | 5 (5-6) | 6 (5-8) | 0.1455 |
| *Baseline blood glucose, median (IQR), mmol/L* | 7.33 (6.83-7.78) | 7.10 (6.70-7.89) | 7.49 (6.95-7.74) | 0.9466 |
| ***TOAST classification*** | | | | |
| *Large-artery atherosclerosis (LAA), n (%)* | 19 (43.2) | 10 (43.5) | 9 (42.9) | >0.9999 |
| *Cardioembolism (CE), n (%)* | 8 (18.2) | 5 (21.7) | 3 (14.3) | 0.7010 |
| *Small-artery occlusion (SAO), n (%)* | 13 (29.5) | 6 (26.08) | 7 (33.3) | 0.7438 |
| *Undetermined etiology, n (%)* | 4 (9.1) | 2 (8.7) | 2 (9.5) | >0.9999 |
| ***Treatment variables*** | | | | |
| *Time from onset to thrombolysis (DNT), median (IQR), min* | 30.5 (27-35) | 30 (26-35) | 31 (29-37) | 0.5624 |
| *Intravenous thrombolysis only, n (%)* | 42 (95.5) | 22 (95.7) | 20 (95.2) | >0.9999 |
| *Bridging mechanical thrombectomy, n (%)* | 2 (4.5) | 1 (4.3) | 1 (4.8) | >0.9999 |

**Table 1. Baseline characteristics**

Data are n (%) or median (IQR). Baseline age, medical history, and pre-thrombolysis NIHSS scores are presented for the overall cohort (n = 44), saline control group (n

= 23), and fructose adjunctive treatment group (n = 21). Continuous variables are shown as median (interquartile range), and categorical variables are presented as number (percentage).

Continuous data were compared using unpaired t-tests, and categorical variables with Fisher’s exact test.

|  | ***Overall***  ***cohort (n=44)*** | ***Control group (n=23)*** | ***Fructose group (n=21)*** | ***P-value*** |
| --- | --- | --- | --- | --- |
| ***Any serious adverse event, n (%)*** | 9 (20.5%) | 5 (21.7) | 4 (19.0) | >0.9999 |
| ***Symptomatic intracranial haemorrhage (sICH) (within 72 hours after thrombolysis), n (%)*** | 1 (2.3%) | 1 (4.3) | 0 (0.0) | >0.9999 |
| ***Systemic haemorrhagic events (within 72 hours after thrombolysis), n (%)*** | 0 (0.0) | 0 (0.0) | 0 (0.0) | / |
| ***Hepatic toxicity (within 30 days post-thrombolysis), n (%)*** | 0 (0.0) | 0 (0.0) | 0 (0.0) | / |
| ***Glycaemic metabolic effects (within 30 days post-thrombolysis), n (%)*** | 8 (18.2) | 4 (17.4) | 4 (19.0) | >0.9999 |

**Table 2. Serious adverse events**

Data are n (%). The safety analysis included all 44 enrolled participants, with 9 serious adverse events recorded overall. Symptomatic intracranial hemorrhage (sICH) was confirmed by cranial CT, accompanied by ≥4-point NIHSS elevation or death, according to SITS‑MOST criteria. Systemic hemorrhagic events conformed to ISTH major bleeding criteria: hemoglobin reduction ≥2 g/dL or transfusion of ≥2 units packed red blood cells. Hemorrhagic endpoints were assessed within 24 hours after thrombolysis. Hepatic toxicity was defined as ALT or AST exceeding 3 times the upper limit of normal (ULN), monitored within 7 days post-thrombolysis. Glycemic metabolic abnormalities referred to newly developed fasting plasma glucose ≥7 mmol/L or HbA1c ≥6.5% within 30 days after thrombolysis. Blood glucose was tested at identical predefined time points in both groups: immediately after infusion completion, the subsequent fasting morning, routine inpatient laboratory screening, and the 30-day follow-up visit to avoid differential detection bias. Between-group comparisons were performed using Fisher’s exact test; P values are denoted as NA when event rates were zero across both groups, as statistical testing was not feasible.

Categorical variables were compared using Fisher’s exact test.

| ***Prognostic indicator*** | ***Overall cohort (n=44)*** | ***Control group (n=23)*** | ***Fructose group (n=21)*** | ***P-value*** |
| --- | --- | --- | --- | --- |
| ***30-day mRS score, median (IQR)*** | 1 (0-1) | 1 (0-1) | 1 (0-2) | 0.9118 |
| ***mRS 0, n (%)*** | 19 (43.2) | 10 (43.5) | 9 (42.9) | >0.9999 |
| ***mRS 1, n (%)*** | 17 (38.6) | 10 (43.5) | 7 (33.3) | >0.9999 |
| ***mRS 2, n (%)*** | 3 (6.8) | 0 (0) | 3 (14.3) | 0.1004 |
| ***mRS 3, n (%)*** | 4 (9.1) | 2 (8.7) | 2 (9.5) | >0.9999 |
| ***mRS 4, n (%)*** | 1 (2.3) | 1 (4.3) | 0 (0) | >0.9999 |
| ***Favorable functional outcome (0-2), n (%)*** | 39 (88.6) | 20 (87.0) | 19 (90.5) | >0.9999 |

**Table 3. Outcomes**

Data are n (%), or median (IQR). Data on neurological function and 30-day functional recovery are summarized across the full cohort (n = 44), control group (n = 23), and fructose adjunctive treatment group (n = 21). Continuous endpoints including Favorable functional outcome and 30-day mRS scores are displayed as median (interquartile range). Categorical mRS stratifications and the proportion of patients with favorable functional outcomes (mRS 0–2) are reported as number (percentage).

Continuous data were compared using unpaired t-tests, and categorical variables with Fisher’s exact test.
